# SMCHD1 loss re-wires MYOD1 enhancer nexuses and chromatin accessibility landscapes in muscle cells

**DOI:** 10.64898/2026.02.21.707202

**Authors:** Zhijun Huang, Wei Cui, Adam Klaiss, Gerd P. Pfeifer

## Abstract

Human SMCHD1 (Structural Maintenance of Chromosomes Flexible Hinge Domain Containing 1) is a chromatin architectural protein linked to heterochromatin repression. Loss of function mutations of SMCHD1 cause facioscapulohumeral muscular dystrophy type 2 (FSHD2) through activation of the *DUX4* homeobox transcription factor gene. However, it is unknown how SMCHD1 may regulate myogenic transcription independently of DUX4. Here, we show that SMCHD1 safeguards enhancer organization within the three-dimensional (3D) genome in human myoblasts. Loss of SMCHD1 leads to widespread gains in chromatin accessibility, aberrant transcription and a global redistribution of the myogenic transcription factor MYOD1. Integrative analyses of histone modifications, chromatin accessibility, Hi-C looping, and activity-by-contact enhancer–gene modeling reveal that SMCHD1 loss rewires the landscape of clustered enhancers and promotes the emergence of a new MYOD1-related network of enhancer elements, termed MYOD1 enhancer nexuses. These structures are marked by increased enhancer–enhancer connectivity, increased local 3D chromatin interactions, and coordinated activation of genes likely relevant for FSHD pathology. Together, our findings identify SMCHD1 as a key architectural constraint that suppresses hyperactive enhancer networks, thereby preserving transcriptional homeostasis in myoblasts.

## INTRODUCTION

The three-dimensional (3D) chromatin architecture serves a fundamental regulatory function that involves transcription factor binding, enhancer activity, and long-range communication between enhancers and genes^1-4^. Therefore, perturbations of higher-order chromatin regulators can lead to widespread transcriptional misregulation^5,6^, even without direct disruption of gene-specific regulatory elements. Understanding how architectural chromatin proteins preserve regulatory stability in lineage-committed cells remains a central challenge in understanding the mechanisms of genome regulation.

SMCHD1 (Structural maintenance of chromosomes flexible hinge domain containing 1) is a noncanonical member of the SMC protein family^7^, with established roles in long-range chromatin repression^8-11^. SMCHD1 plays important roles in X-chromosome inactivation^7-9^, genomic imprinting^12-14^, and repression of repetitive or development-associated loci^13,15-17^, suggesting that the primary function of SMCHD1 is shaping chromatin topology rather than functioning as a conventional transcriptional regulator. Moreover, in muscle cells, aberrant functioning of SMCHD1 is associated with transcriptional dysregulation and disease phenotypes^18-22^. However, the genome-wide architectural mechanisms by which SMCHD1 organizes regulatory chromatin in myogenic cells remain not fully defined.

Skeletal myogenesis is orchestrated by the specialized hierarchy of lineage-determining transcription factors^23,24^, among which MYOD1 functions as a central regulator of enhancer activation and chromatin remodeling^25,26^. MYOD1 binding is highly influenced by chromatin accessibility and local genomic architecture^27,28^. Furthermore, MYOD1 binds to large regulatory domains with clusters of enhancers, which are associated with genes that predominantly contribute to the identity of the myogenic cell^29^. These super-enhancers associated with MYOD regulate cell identity by rewiring the three-dimensional genome organization^29,30^. These enhancer clusters function as regulatory domains that control long-range enhancer–gene interactions to drive muscle-specific gene expression programs. However, whether SMCHD1 directly orchestrates lineage-defining transcription factor–driven enhancer networks and higher-order enhancer–gene connectivity in myogenic cells has not been established.

Recent methodological advances enable systematic analysis of enhancer activity within a 3D genomic context. Activity-by-Contact (ABC) modeling integrates analyzing chromatin contacts, enhancer activity, and physical proximity to estimate functional enhancer– promoter communication^31^. In addition, high-resolution Hi-C and loop-based analysis provide more detailed insights into the landscape of chromatin interactions. Together, these approaches offer a powerful framework to dissect how chromatin architecture regulates enhancer-driven transcriptional programs.

A hallmark of FSHD is the abnormal activation of the embryonic transcription factor gene *DUX4* in diseased muscle tissue^32-35^. Here, we applied integration analysis of a multi-omic strategy combining RNA-seq, ATAC-seq, ChIP-seq, Hi-C looping, and ABC-based enhancer– gene modeling to define the function of SMCHD1 in human myoblasts. We demonstrate that the loss of SMCHD1 results in widespread transcriptional dysregulation independent of *DUX4* activation, accompanied by global increases in chromatin accessibility and perturbation of the 3D genome landscape. SMCHD1 depletion leads to a pronounced redistribution of MYOD1 binding across the genome and a reconfiguration of the super-enhancer landscape. Strikingly, SMCHD1 loss promotes the emergence of new MYOD1-associated enhancer networks - here termed MYOD1 enhancer nexuses - characterized by coordinated enhancer activation, increased local chromatin accessibility, and strengthened enhancer–gene looping. These hyper-interactive enhancer clusters arise preferentially from super-enhancers and stitched enhancers and drive robust transcriptional activation of myogenic and disease-relevant gene programs in the absence of SMCHD1. Collectively, our findings demonstrate that SMCHD1 is a key chromatin architectural regulator that limits enhancer network formation and preserves transcriptional stability by modulating MYOD1-related enhancer networks in human myoblasts.

## RESULTS

### Loss of SMCHD1 leads to aberrant expression of genes independent of DUX4

We previously engineered the male immortalized human myoblast cell line LHCN-M2 with a CRISPR knockout strategy to obtain stable SMCHD1-deletion cell lines^10^, and identified 388 differentially expressed genes (fold change > 2, FDR < 0.05) with mRNA seq^10^. To identify the full transcriptomic changes induced by SMCHD1 loss, we generated total RNA-seq data from wild-type and SMCHD1-deficient cells. We found 271 upregulated genes and 227 downregulated genes (fold change >2, FDR<0.05), (Fig. 1A; Supplementary Table 1). Notably, analysis of the DAVID^36-38^ GO terms showed enrichment of terms related to muscle cell development (Fig. 1B). The difficulty of detecting *DUX4* transcripts in FSHD2 patient samples has been reported by several independent studies^35,39-41^, indicating that SMCHD1 defects might promote an additional pathological function independent of DUX4. We evaluated the DUX4-related specific marker genes reported in FSHD2^40^ and found that SMCHD1 inactivation did not significantly change the expression of the FSHD2-specific markers (Fig. 1C). By examining known DUX4 downstream-regulated genes in human myoblasts^42^, only a small number of genes was affected by SMCHD1 depletion; notable among these downregulated targets was the myogenic regulatory factor, *MYF5* (Fig. 1D). The other changes were not significant. Indeed, we did not observe DUX4 expression in the wildtype myoblasts and any activation by the loss of SMCHD1 (Fig. 1E). Thus, the differentially expressed genes observed in the absence of SMCHD1 were not related to DUX4 expression. In addition to the large number of differentially expressed genes we found in the previous mRNA-seq analysis^10^, loss of SMCHD1 provoked the differential expression of 46 different non-coding RNAs (Fig. S1A), 3 miRNA, 3 snRNA, 6 antisense genes, 13 LincRNAs and 10 pseudogenes (Fig. S1B). Gene functional enrichment analysis showed that the differentially expressed LincRNAs after SMCHD1 deletion were related to muscle tissue development, among several other terms (Fig. S1C). Therefore, the alteration of the transcriptome caused by SMCHD1 deletion is related to muscle development, and the change in function was independent of DUX4. We did not observe that the cell cycle (Fig. S2A) and cell proliferation abilities (Fig. S2B) of myoblasts were affected by the absence of SMCHD1. These results suggest that SMCHD1 is involved in the progression of muscle development, while its downregulation might trigger mechanisms independent of DUX4.

**Figure 1:**
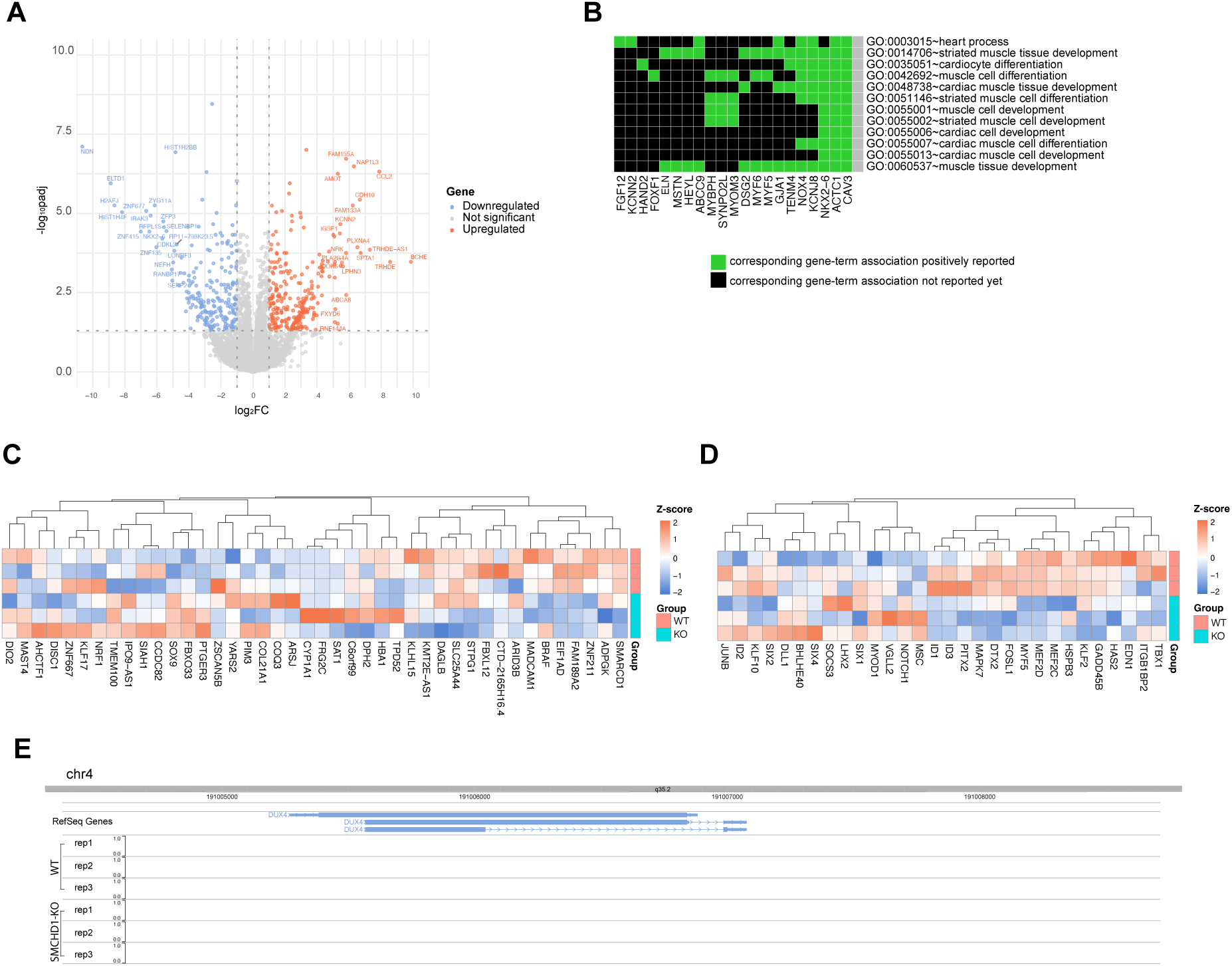
SMCHD1 loss leads to aberrant expression of genes in myoblasts, independent of DUX4 target gene activation. **A.** Volcano plot showing the differentially expressed genes (DEGs) after SMCHD1 knockout. All DEGs with FDR <=0.05 and log 2-fold change >2 are labeled in light blue and red. **B.** Heatmap showing the DAVID functional annotation of differentially expressed genes. **C.** Heatmap showing the expression of genes differentially expressed in the DUX4-affected cells of the FSHD2 samples identified by Silvère van der Maarel et. al.^40^ upon SMCHD1 knockout. **D.** Heatmap showing the expression of DUX4 target genes identified by Michael Kyba et. al.^42^. **E.** WASH U browser showing the RNA expression of *DUX4* in wildtype and SMCHD1 knockout LHCN-M2 cells.

### SMCHD1 governs the chromatin accessibility landscape of myoblasts

Since one of the consequences of SMCHD1 loss is characterized by altered chromatin domains^9-11,15,43^, we then asked how SMCHD1 governs chromatin status. To answer this question, we used ATAC-seq to detect transposase-accessible chromatin in WT and SMCHD1-knockout LHCN-M2 cells. In total, 21276 differential accessible regions (DARs) were identified. We found 12640 increased accessible regions (59.4%) and 8636 reduced accessible regions (40.6%) (Fig. 2A). The functional enrichment analysis of the DAR-related genes showed that both increased (Fig. 2B) and reduced accessible regions (Fig. 2C) after SMCHD1 loss were mainly related to neuron development. Muscle tissue development was also an enriched term for both upregulated and downregulated DARs. Combining the ATAC-seq data with the RNA-seq data, our analysis revealed that the promoters of up-regulated differentially expressed genes (DEGs) were characterized by increased chromatin accessibility (Figure 2D-2E), whereas down-regulated DEGs were associated with more condensed chromatin states (Figure 2D-2E). When placing these genes in the context of muscle development, particularly inflammatory response to muscle injury-related genes, such as *CCL2*^44,45^, and muscle development and function-associated genes, such as *SELENBP1*^46,47^, they typically exhibited increased chromatin accessibility at the promoter regions or suppressed chromatin states following SMCHD1 depletion, respectively (Figure 2E).

**Figure 2:**
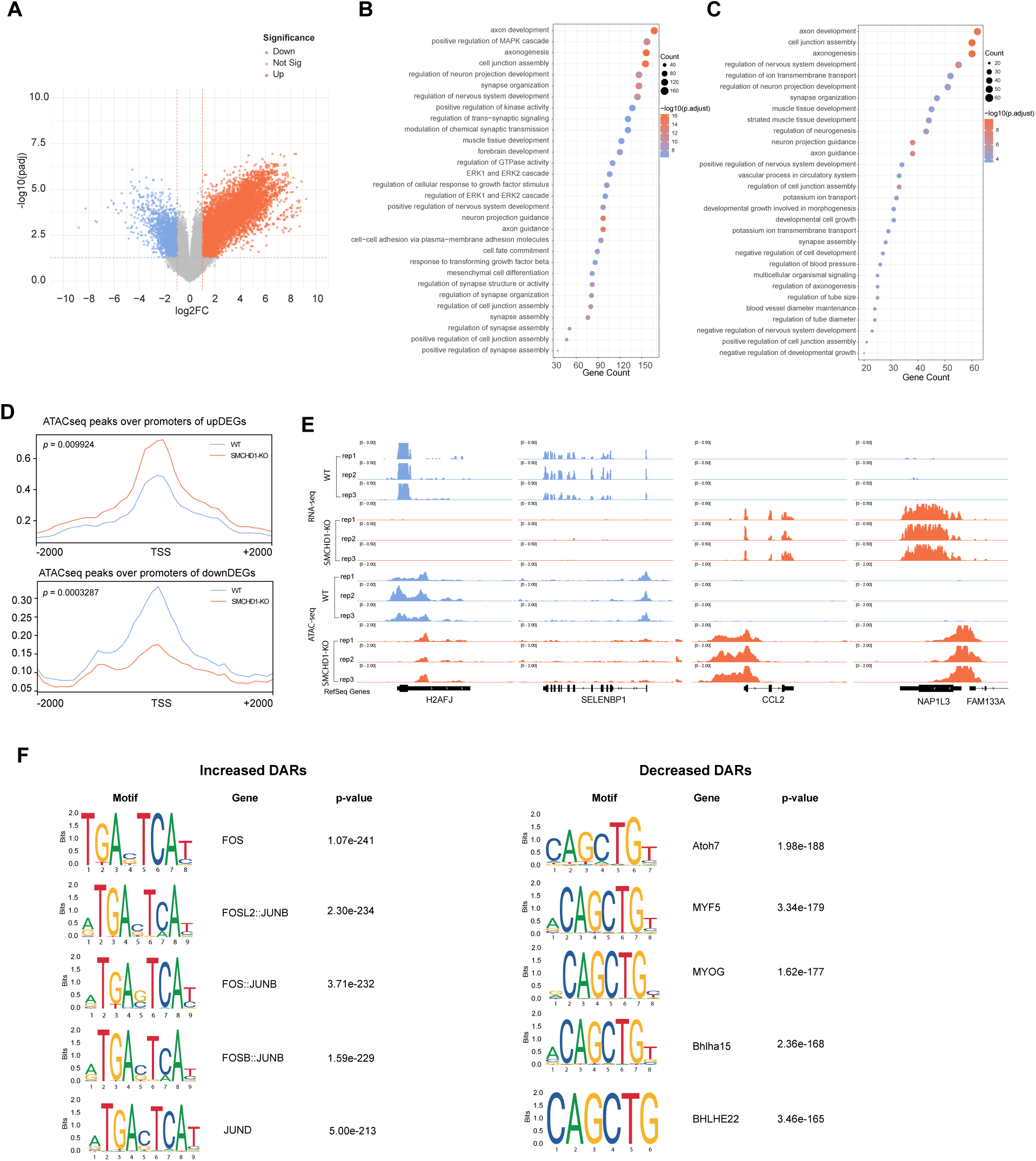
SMCHD1 loss alters the chromatin accessibility landscape of myoblasts. **A.** Volcano map showing the DARs (Differentially Accessible Regions) in LHCN-M2 cells after SMCHD1 knockout. DARs detected by ATAC-seq with FDR <= 0.05 and log2 fold change >2 were labeled in light blue and red, respectively. **B.** Gene functional analysis of increased DARs upon SMCHD1 knockout. **C.** Gene functional analysis of decreased DARs upon SMCHD1 knockout. **D.** Profile plots of ATAC-seq showing increased chromatin accessibility at the promoters of upregulated genes, and reduced chromatin accessibility at the promoters of down-regulated genes caused by SMCHD1 depletion. A paired t-test was used for statistical analysis for calculating chromatin accessibility differences. **E.** IGV browser views of ATAC-seq peaks and RNA-seq peaks at the promoter regions of genes showing down-regulation (*H2AFJ* and *SELENBP1*) and up-regulation (*CCL2* and *NAP1L3*). **F.** Motif analysis of increased and reduced DARs.

To determine whether motifs for specific transcription factors are associated with DARs in the absence of SMCHD1, we performed motif analysis for the increased and reduced DARs. The increased DARs were enriched for the JUN/FOS like motif “TGAcTCA”, and motif “CAGCTG” for the bHLH family of transcription factors showed the top propensity for decreased DARs (Fig. 2F). The MYF5-like motif enriched in decreased DARs was consistent with the downregulation of *MYF5* in the SMCHD1 knockouts (Fig. S3B). Interestingly, the JUN-like motif was enriched in increased DARs but without any aberrant gene expression change of the *JUN/FOS* family (Fig. S3C, S3D). Genome-wide MYOD1 binding occurs in chromatin contexts that are permissive for AP-1 (JUN/FOS) factor binding, implying functional interplay between MYOD1 and JUN-associated regulatory programs^48,49^, and suggesting the potential function of MYOD1 in orchestrating chromatin accessibility and 3D genome organization in the absence of SMCHD1. Although we did not observe a significant transcriptional change of *MYOD1* (Fig. S3A), the redistribution of MYOD1 after SMCHD1 inactivation would be a reasonable possibility to be considered.

### SMCHD1 safeguards chromatin accessibility and 3D-genome architecture

The Hi-C-based 3D structure of the human myoblast genome was partitioned into either the active “A” compartments or the inactive “B’ compartments based on our previous analysis^10^. Our data showed that the newly formed B-to-A compartment transition regions were highly enriched in open chromatin signals in the absence of SMCHD1, whereas A-to-B compartment regions were mainly enriched in condensed chromatin signals, as determined by calculating the number (Fig. 3A) and intensity (Fig. 3B) of the DARs in the compartment switching regions. We observed that loss of SMCHD1 promoted B-to-A compartment transitions, which were associated with more open chromatin signals (Fig. 3C, Fig. S4A). When zooming in to reveal further details, many new topologically associated domains (TADs) emerge in parallel with the B-to-A compartment transitions^10^. Increased ATAC-seq chromatin signals were enriched in the newly formed TAD regions (Fig. 3D). Increased chromatin accessibility and new TADs at representative chromosomes are shown as examples (Fig. 3D, Fig. S4B-S4C). Globally, chromatin accessibility of all newly formed TADs in B-to-A switched regions was dramatically increased by SMCHD1 deletion (Fig. 3E), whereas it was decreased in the WT samples. Our further comparisons revealed increased contacts in chromatin loop anchors in B-to-A regions in the absence of SMCHD1 (Fig. 3F-H). Moreover, the increased chromatin accessibility at the loop anchors in B-to-A regions was seen at the whole-genome level (Fig. 3I). In situ Hi-C contact maps combined with ATAC-seq tracks are depicted in Figure 3J and showed gains of loops at representative regions on chromosome 4, linked to strong chromatin accessibility in SMCHD1 KO cells (Fig. 3J), and a similar pattern at additional representative regions is shown in Figure S4D. These results further support the role of SMCHD1 in mediating the 3D structure and chromatin accessibility of the human muscle cell genome.

**Figure 3:**
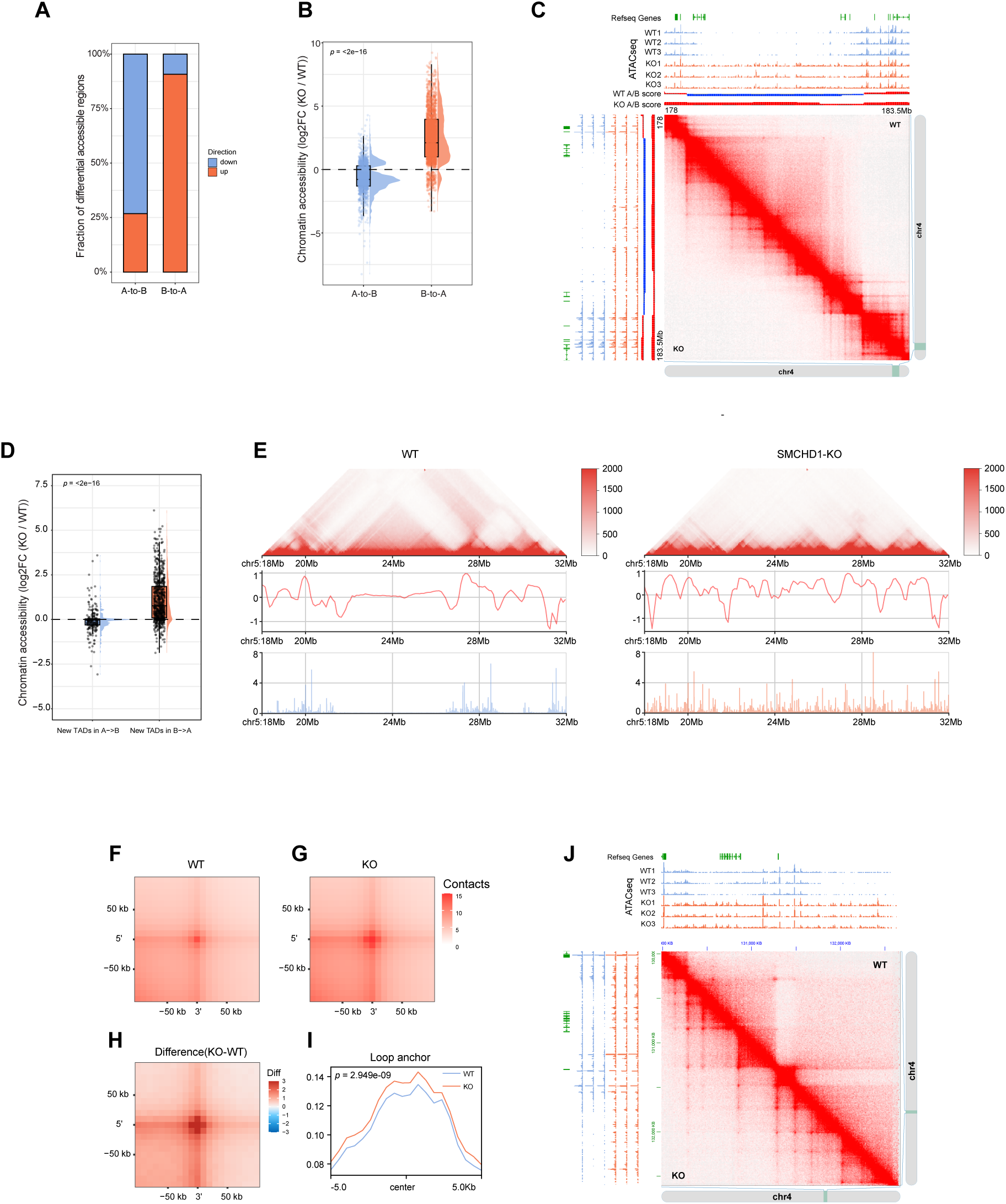
SMCHD1 safeguards the chromatin accessibility and nuclear architecture of myoblasts. **A.** Stacked bar chart showing the fraction of DARs (up or down) in compartment switching regions. **B.** Rain-cloud plot showing the log2 fold change of ATAC-seq signal in compartment switching regions. The midline is the median of the data, with the upper and lower limits of the box being the third and first quartile (75th and 25th percentile), respectively. Whiskers extend to the most extreme data points within 1.5 × inter-quartile range (IQR) from the box bounds, and minima and maxima within the whiskers indicate the range of non-outlier data. Statistical significance was assessed using a two-sided Wilcoxon rank-sum test (*p* < 2.2e − 16). **C.** In situ Hi-C contact map showing compartment changes across the 178-183.5 Mb region of chromosome 4. The ATAC-seq signals of WT and SMCHD1 KO are shown along with the contact map on top and left. **D.** Rain-cloud plot showing the log2 fold change of ATAC-seq signal in newly formed TAD boundaries. The midline is the median of the data, with the upper and lower limits of the box being the third and first quartile (75th and 25th percentile), respectively. Whiskers extend to the most extreme data points within 1.5 × inter-quartile range (IQR) from the box bounds, and minima and maxima within the whiskers indicate the range of non-outlier data. Statistical significance was assessed using a two-sided Wilcoxon rank-sum test (*p* < 2.2e − 16). **E.** Hi-C contact map and insulation profiles at a representative region on chromosome 5 in wildtype cells (left panel) and SMCHD1knockout cells (right panel). The ATAC-seq signals of WT and SMCHD1 KO are shown at the bottom. **F-H.** Aggregate peak analysis (APA) of chromatin loops in B-to-A regions showing increased interaction intensity in SMCHD1 knockout cells (G) versus wild-type cells (F). The differential APA analysis plot with red denotes more interactions in SMCHD1 knockout cells compared to the wildtype cells (H). **I.** Profile plot of ATAC-seq signal showing increased chromatin accessibility at the anchor of loops in B-to-A switched regions. A paired t-test was used for statistical analysis for calculating chromatin accessibility differences. **J.** In situ Hi-C contact map showing gains of loops at representative regions on chromosomes 4, linked to newly arising chromatin accessibility in SMCHD1 KO cells as shown in the ATAC-seq tracks along with the contact map on top and left.

### SMCHD1 loss triggers global redistribution of MYOD1 across the genome

To address whether depletion of SMCHD1 drives MYOD1 occupancy changes in human myoblasts, we performed MYOD1 ChIP-seq in wildtype and SMCHD1-knockout human myoblasts. The results showed that 4511 MYOD1-binding regions were identified in the human myoblasts. We found that a significant proportion of MYOD1-occupied peaks were at intergenic regions, introns and promoters (Fig. 4A), consistent with MYOD1 activity involving the regulation of gene expression through both enhancer and promoter regions^50,51^. Further motif analysis revealed that MYOD1-occupied regions are enriched for E-boxes in our datasets (Fig. 4B), confirming previous reports^48,49^. Moreover, profiles of MYOD1 enrichment and chromatin accessibility revealed that MYOD1 showed increased binding at open chromatin regions, while decreased binding at more condensed chromatin regions (Fig. 4C, 4D). Furthermore, we identified 190 increased binding and 137 decreased MYOD1 peaks after SMCHD1 loss (Fig. 4E). We next identified 21 significantly differentially expressed genes (Fold change>2 and FDR < 0.05) that were related to the differential binding of MYOD1 calculating physical proximity of differential binding sites, when SMCHD1 was deleted (Fig. 4F, Fig. S5A). Notable among these upregulated targets were *FAM155A*, *CCL2* and *FRG2B*. *FAM155A* encodes a component of hetero-tetrameric NALCN sodium channels^52-54^. Alterations in the NALCN sodium leak channels, which FAM155A is a component of, have been linked to hypotonia, a form of muscle weakness^55^. *FRG2B* belongs to a gene family specifically upregulated in FSHD myoblasts and has been considered earlier as a key candidate gene in disease pathogenesis^56^. Similarly, the inflammatory chemokine CCL2 is a known regulator of muscle phenotypes and strength, typically increasing in response to myogenic injury^44,45^. The induction of these genes suggested that SMCHD1 deficiency triggers a transcriptional program that may contribute to the FSHD2 disease state. RNA-seq, ATAC-seq, MYOD1 ChIP-seq tracks, and ChromHMM chromatin state maps are depicted in Figure 4H and showed that induction of gene expression is linked to the more open chromatin accessibility signature and more MYOD1 binding at the enhancer or promoter for the *CCL2* gene in the absence of SMCHD1 (Fig. 4G). A similar binding pattern was also observed at the *FRG2B* and the *FAM155A* locus (Fig. S5B-5C). On the other hand, differential MYOD1 binding was also related to down-regulated genes with a different binding pattern (Fig. S5E-5F). Further correlation analysis between differential gene expression and differential MYOD1 binding showed a weak positive correlation (Fig. S5A), suggesting that MYOD1 may be involved in gene regulation not only through its physical proximity but also through long-distance interactions. This led us to examine whether MYOD1 differential binding is preferentially present with a specific chromatin status. Combining ChromHMM of myoblasts with our MYOD1-ChIP seq data, we found that a significant proportion of differential MYOD1 binding peaks were in enhancer regions (Fig. 4H, Fig. S6A-6B), and increased MYOD1 binding is only seen in the heterochromatin regions when SMCHD1 is lost (Fig. 4H). Collectively, these results suggested that the MYOD1 differential binding was related to the 3D structure change caused by SMCHD1 inactivation.

**Figure 4:**
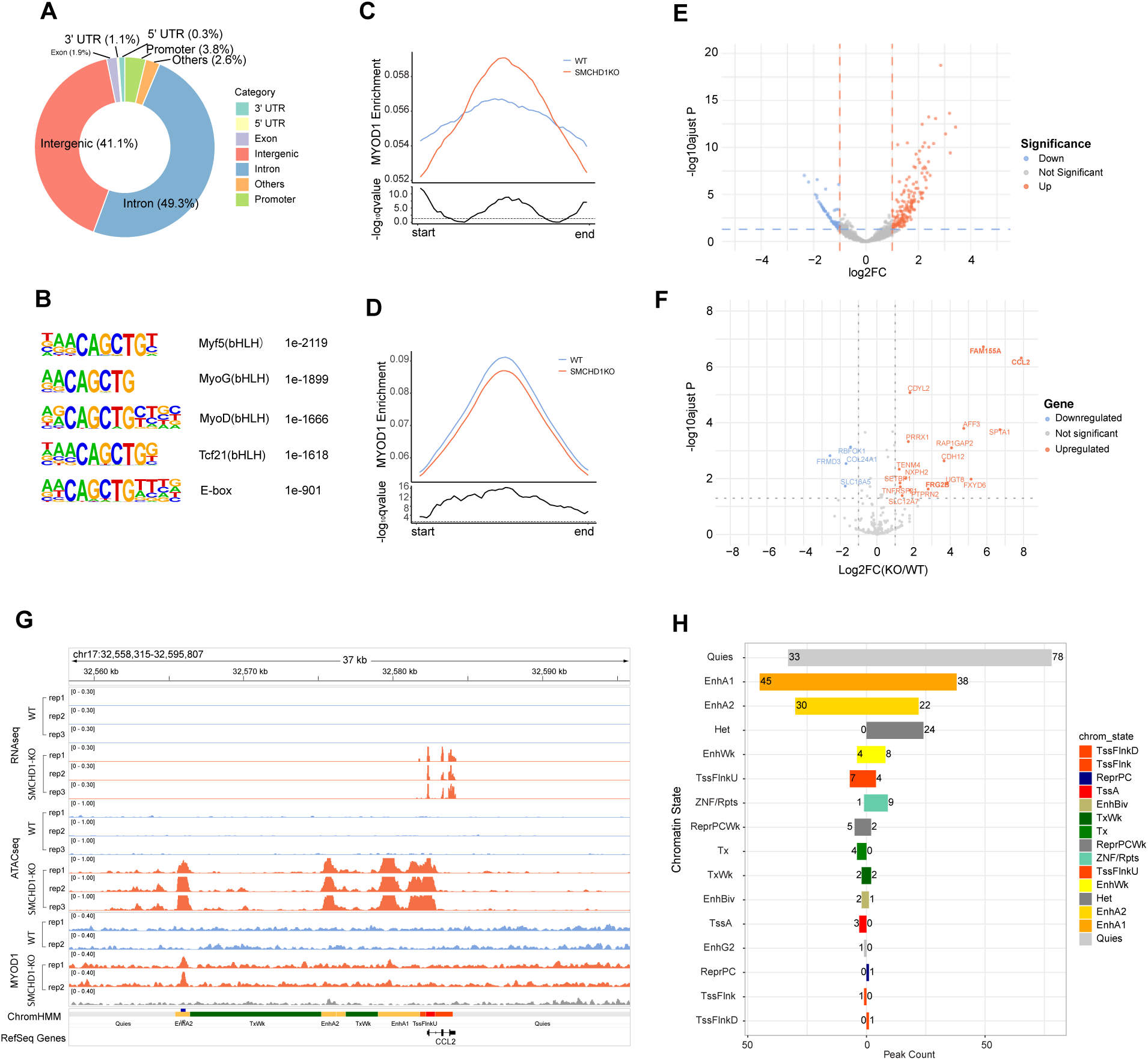
Loss of SMCHD1 reshapes the MYOD1 distribution in the genome. **A.** The donut pie chart shows the fraction of MYOD1 binding sites in genome compartments of wildtype myoblasts. **B.** Motif analysis of MYOD1 binding sites. **C.** Profile plot of MYOD1 ChIP-seq showing increased MYOD1 enrichment at increased chromatin accessibility regions detected by ATAC-seq (upper panel). Significance of the differences (paired t-test), the line above the dash line (*q*-value < 0.05) indicates a statistically significant difference of KO versus WT cells (lower panel). **D.** Profile plot of MYOD1 ChIP-seq showing decreased MYOD1 enrichment at reduced chromatin accessibility regions detected by ATAC-seq (upper panel). Significance of the differences (paired t-test), the line above the dash line (*q*-value < 0.05) indicates a statistically significant difference of KO versus WT cells (lower panel). **E.** Volcano plot showing the differential binding sites of MYOD1 after SMCHD1 knockout. All differential binding sites with FDR <= 0.05 and log2 fold change >2 were labeled in light blue and red. **F.** Volcano plot showing the expression of differential MYOD1 binding genes. Differentially expressed genes with FDR <= 0.05 and log2 fold change >2 were labeled in light blue and red. **G.** Illustration of ATAC-seq peaks, RNA-seq peaks and MYOD1 binding sites at the *CCL2* gene locus. **H.** Bar chart showing the fraction of differential MYOD1 binding sites in different chromatin states.

### SMCHD1 loss rewires the super-enhancer landscape and establishes new MYOD1-enhancer nexuses in human myoblasts

Our ATAC-seq analysis revealed a subset of differential accessibility regions that were not isolated but rather formed dense clusters at enhancers (e.g., Fig. 4G). Moreover, the enhancer regions were enriched with a more open chromatin signature and more specific MYOD1 binding (Fig. S6C-6D). To determine if these regions constitute higher-order regulatory units in the WT and SMCHD1 knockouts, we applied the ROSE algorithm^29^, “stitching” together individual enhancers within 12.5 kb of each other. These stitched regions were then treated as single regulatory units for subsequent H3K27ac and H3K4me1 signal ranking and classification. Both stitched enhancers and the remaining individual enhancers (those lacking a neighbor within 12.5 kb) were included in this ranking. A subset of these elements exhibited exceptionally high levels of H3K27ac or H3K4me1 modifications, defining them as super-enhancers (SE) (Fig. 5A-B, 5D-E), according to the standard ROSE criteria^29^. For downstream analysis in this study, we identified a core set of super-enhancers by overlapping the lists generated independently from the H3K27ac and H3K4me1 rankings. This ensures high confidence in these regulatory regions, suggesting only genomic regions classified as super-enhancers by both H3K27ac and H3K4me1 enrichment criteria were considered bona fide super-enhancers in this study (Fig. 5C, 5F). In total, 977 of SE for WT and 777 of SEs for SMCHD1 knockout myoblast cells were identified at the genome level (Fig. 5C, 5F and Fig. S7). The number of SEs in SMCHD1 knockout is smaller than the WT, because more H3K27ac and H3K4me1 binding caused by SMCHD1 deletion results in denser clusters being designated as a regulatory unit, which is in accordance with denser H3K27ac and H3K4me1 binding regions being identified by SMCHD1 loss (Fig. S7).

**Figure 5:**
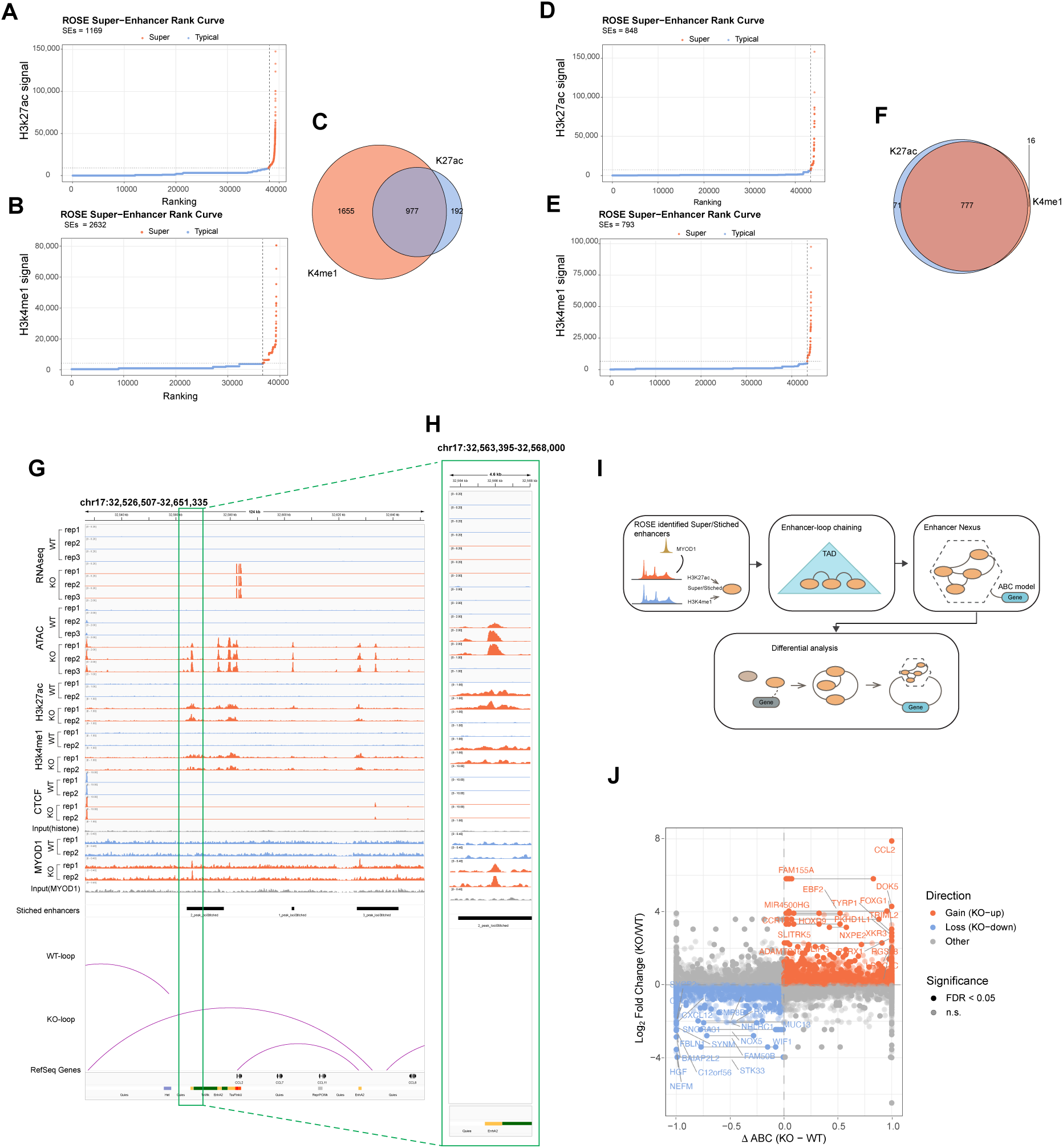
SMCHD1 loss reorganizes the super-enhancer landscape and establishes MYOD1 mediated regulatory networks in myoblasts. **A-B.** Identification of super enhancers in wild-type LHCN-M2 myoblast. H3K27ac (**A**) and H3K4me1 (**B**) signal (as defined by the ROSE ranking algorithm) enrichment represented for each enhancer region. **C.** Super enhancer overlap of high H3K27ac and H3K4me1 enrichment for wild type cells. **D-E.** Identification of super enhancers in SMCHD1 knockout LHCN-M2 myoblasts. H3K27ac (**D**) and H3K4me1 (**E**) signal (as defined by the ROSE algorithm) enrichment represented for each enhancer region. **F.** Super enhancer overlap of high H3K27ac and H3K4me1 enrichment for SMCHD1 knockout cells. **G-H.** Illustration of the *CCL2* gene in IGV browser views showing gene expression related to the more open chromatin accessibility, H3K27ac and H3K4me1 binding, and more loop contacts. The stitched enhancers called by ROSE were bound by newly formed MYOD1 upon SMCHD1 deletion (**G**). Enlargement of an area at the gene locus from panel G (**H**). **I.** MYOD1 enhancer nexus identification summary. **J.** Volcano-like scatter plot displaying the relationship between MYOD1-related enhancer cluster-gene contact activity (ΔABC, KO – WT, x-axis) and gene expression changes (log₂FC, KO/ WT, y-axis) across all enhancer cluster–gene pairs. Each point represents a gene linked to at least one enhancer cluster by the ABC model. Points are colored by the direction of transcriptional change: KO-upregulated genes (red), KO-downregulated genes (blue), and nonsignificant or unchanged genes (gray). Circle fill indicates statistical significance (solid represents FDR < 0.05, half-solid represents not significant).

To investigate the regulatory mechanisms by which SMCHD1 loss induces gene expression, we initially employed the standard ROSE algorithm to identify potential super-enhancer-associated genes. This method assigns regulatory targets based on linear genomic proximity, including the nearest or overlapping genes. Surprisingly, we found that simple physical proximity was not a perfect predictor of transcriptional output, as many “related” genes showed no significant change in expression despite their proximity to a super-enhancer (Fig. S8). This observation was consistent with recent findings that linear distance often fails to capture functional enhancer-promoter interactions. Previous studies have demonstrated that 3D chromatin contacts, rather than linear genomic distance, can provide a more accurate and biologically relevant framework for identifying true regulatory targets^57,58^. For instance, integrating 3D contact frequency significantly improves the identification of functional interactions^31,59^. Therefore, given SMCHD1’s role in chromatin structure, we further investigated whether MYOD1 and these super-enhancers direct specific gene expression programs in the absence of SMCHD1. Using a “loop-chaining” approach, a custom R script, we mapped high-confidence spatial interactions to identify a core subset of MYOD1-enhancer nexuses that overcome the limitations of 1D genomic distance (Fig. 5I). Here, we defined MYOD1-enhancer nexuses by integrating MYOD1 ChIP-seq, active enhancer chromatin marks, HiC–derived chromatin interactions, and a modified ABC-based enhancer–gene linking model (Fig. 5I). MYOD1-related clustered enhancers were physically connected through chained chromatin loops and were grouped into higher-order regulatory units, termed the MYOD1 enhancer nexuses, which localized within single TADs and displayed dense intra-nexus connectivity. Loss of SMCHD1 led to pronounced activation of these nexuses, marked by increased chromatin accessibility, strengthened chromatin contact looping, and elevated ΔABC scores. Nexus activation was tightly coupled to robust upregulation of linked target genes, indicating that SMCHD1 normally restrains MYOD1-involved enhancer network formation and transcriptional outputs (Fig. 5I). We found that SMCHD1 loss resulted in active MYOD1-enhancer nexuses, which was associated with upregulated expression of specific genes (Fig. 5J). Notably, the induction of *FAM155A* and *CCL2* could be explained by this model (Fig. 5G, Fig. 5H, Fig. S9A).

### SMCHD1 loss activates MYOD1 enhancer nexus programs through enhancer rewiring and 3D chromatin looping

To systematically characterize how SMCHD1 loss reshapes the MYOD1-associated regulatory architecture, our study combined condition-specific ABC modeling, loop– chaining–based enhancer nexus identification, chromatin accessibility, and transcriptional profiling. It enables a quantitative assessment of enhancer-gene communication rewiring and its transcriptional consequences following SMCHD1 inactivation. We identified MYOD1 enhancer nexuses and revealed widespread enhancer activity in the nexus (Fig. 6A). Moreover, both super-enhancers and stitched enhancers contributed to activity gains of the enhancer nexus (Fig. 6A), indicating coordinated activation of large enhancer assemblies rather than isolated regulatory elements. Notably, MYOD1-enhancer nexuses ranked by the mean transcriptional response of linked genes showed a clear correspondence between elevated ΔABC values and transcriptional induction (Fig. 6B), suggesting functional coupling between enhancer rewiring and gene activation in the absence of SMCHD1.

**Figure 6:**
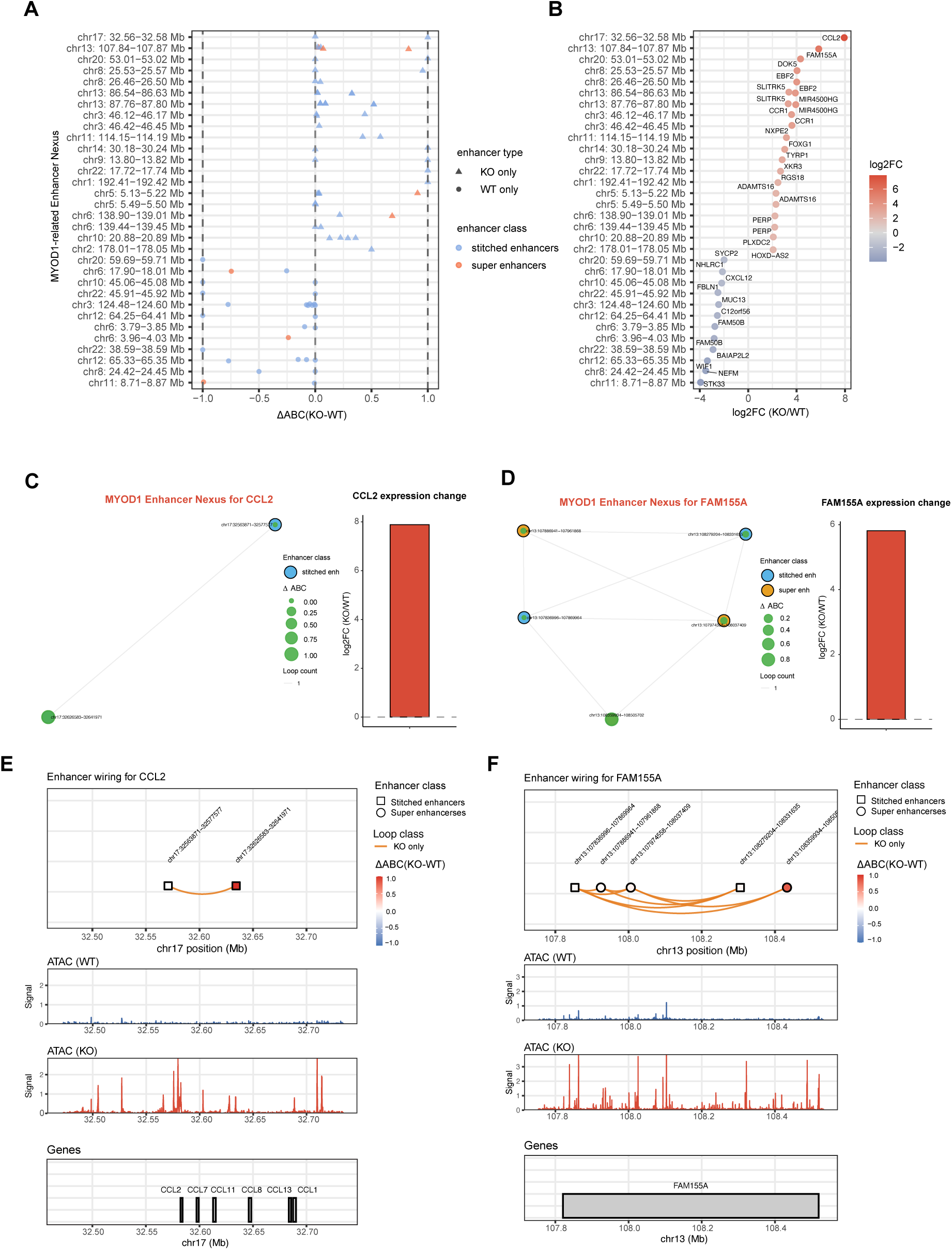
SMCHD1 loss induces MYOD1 enhancer nexus activation and associated transcriptional responses by altering chromatin accessibility, enhancer–gene wiring, and 3D chromatin looping. **A.** ΔABC scatterplot showing super and stitched enhancer-level activity changes across MYOD1 enhancer nexuses. Each row represents a MYOD1 enhancer nexus, ordered top-to-bottom by the mean log₂FC of significantly regulated MYOD1 enhancer nexus-linked genes (panel B). Individual points correspond to enhancers within each MYOD1 enhancer nexus. Point shape indicates enhancer class specificity derived from enhancer presence or absence between WT and SMCHD1-KO. Point color distinguishes enhancer categories: orange indicates super-enhancers; blue indicates stitched enhancers. The X-axis shows the ΔABC (KO − WT) enhancer activity derived from condition-specific ABC model calculations. Dashed vertical lines at ΔABC = −1 and +1 denote thresholds illustrating high-activity enhancer rewiring events. **B.** MYOD1 enhancer nexus -linked significant genes identified using condition-specific ABC model calculations. For each enhancer nexus in (**A**), its related significantly dysregulated genes (FDR < 0.05 and |log₂FC| > 2) are plotted according to log₂FC of RNAseq. Blue and red points indicate downregulated and upregulated expression, respectively. Rows are aligned identically between panel A and panel B. **C.** MYOD1 enhancer nexus activated by SMCHD1 loss for *CCL2*. Two KO-only enhancer clusters form a MYOD1 enhancer nexus upstream of *CCL2*, each showing strong enhancer activity, and KO-induced looping. Node size indicates the enhancer ABC activity, *CCL2* expression exhibits a strong KO-induced increase, as shown in the bar chart at right. **D.** MYOD1 enhancer nexus activated by SMCHD1 loss for *FAM155A*. Five KO-only stitched and super-enhancers converge on *FAM155A*, forming a robust KO-specific MYOD1 enhancer nexus with high enhancer activity and strong KO-upregulation of the target gene. Node size indicates enhancer ABC activity, *FAM155A* expression exhibits a strong KO-induced increase, as shown in the bar chart at right. **E-F.** Enhancer wiring, peak tracks, and gene models for the representative MYOD1 enhancer nexuses for *CCL2* and *FAM155A*. Top: genomic enhancer wiring tracks show enhancer coordinates (black squares), KO-only enhancer arcs (orange), and ΔABC activity is defined by arc color. Middle: ATAC-seq signal confirms KO-specific chromatin opening at enhancers. Bottom: gene tracks illustrate the position of the gene relative to the enhancer clusters in the nexus.

To illustrate the structural and functional organization of a MYOD1 enhancer nexus, we examined representative loci activated upon SMCHD1 loss. At the *CCL2* locus, we observed that two KO-only enhancer clusters formed a SMCHD1-dependent MYOD1 enhancer nexus characterized by high ABC activity and strong KO-specific looping interactions (Fig. 6C). Both enhancer clusters exhibited increased chromatin accessibility and enhancer activity in knockout cells, accompanied by a significant upregulation of *CCL2* expression (Fig. 6C). Another representative nexus architecture was observed at the *FAM155A* locus, where multiple KO-only stitched and super-enhancers formed a robust MYOD1 enhancer nexus (Fig. 6D). These enhancers displayed high ABC activity and extensive KO-specific looping, collectively driving strong transcriptional activation of *FAM155A*. Together, this network visualization highlighted strengthened enhancer–gene connections, indicating that enhancer activation and loop reinforcement occur concomitantly following SMCHD1 depletion. The nexus model was further supported by the integration of enhancer wiring diagrams with epigenomic profiles (Fig. 6E–F). For representative loci, *CCL2* and *FAM155A*, KO-specific enhancer arcs corresponded to regions of increased chromatin accessibility signals (Fig. 6E–F). Although we found that some WT-only nexuses were found in WT samples, which were associated with more transcriptional activity for specific genes in WT, the chromatin accessibility and contacts were not decreased in the SMCHD1 knockout (Fig. S9B–9E), suggesting that the decreased gene expression is not caused by the spatial structural changes of chromatin at the nexus. These changes indicated that SMCHD1 loss promotes chromatin opening at enhancer elements, facilitates enhancer–gene communication through enhanced looping, and enables the formation of the MYOD1 enhancer nexus that drives transcriptional activation of target genes in human myoblasts.

### Loss of SMCHD1 increases 3D contacts and local chromatin accessibility at MYOD1 enhancer nexuses

To further elucidate whether MYOD1 enhancer nexus activation upon SMCHD1 loss is accompanied by changes in local chromatin architecture, we examined chromatin accessibility and 3D contact organization at enhancer elements within MYOD1 enhancer nexuses. ATAC-seq profiling revealed a significant global increase in chromatin accessibility across all enhancer elements within the MYOD1 enhancer nexuses in SMCHD1-KO cells (Fig. 7A), indicating that SMCHD1 loss broadly promotes chromatin opening at MYOD1-associated enhancers. To determine if the increased enhancer accessibility was accompanied by alterations in local 3D chromatin interactions, we utilized GENOVA^60^ computed chromatin contact frequencies centered on enhancer elements within the MYOD1 enhancer nexus. WT cells exhibited a contact pattern around enhancer elements that was relatively weak and diffuse, suggesting limited local interaction density (Fig. 7B, left). In contrast, SMCHD1-knockout cells displayed a pronounced increase in contact intensity centered on enhancer elements (Fig. 7B, middle). Difference heatmaps (KO − WT) revealed widespread gains in contact frequency within approximately ±200 kb of enhancer elements, indicating enhanced local chromatin interactions following SMCHD1 loss (Fig. 7B, right). The chromatin structural consequences of SMCHD1 depletion were further confirmed by 3D surface representations of these contact matrices (Fig. 7C). WT cells exhibited a relatively flat interaction landscape over super-enhancer and stitched enhancer regions, while SMCHD1-knockout cells showed prominent local peaks, reflecting increased contact density and enhanced self-interaction of clustered enhancers in the nexus. These data lead to a model in which SMCHD1 governs the MYOD1 enhancer nexus landscape by limiting excessive chromatin accessibility and maintaining a repressive chromatin environment at latent enhancers. SMCHD1 loss not only increases chromatin accessibility but also strengthens local 3D chromatin interactions around enhancer elements at MYOD1 enhancer nexus positions. Together, the loss of these structural constraints allows for the formation of a hyper-interactive enhancer nexuses that drives the myogenic and FSHD-related transcriptional programs (Fig. 7D).

**Figure 7:**
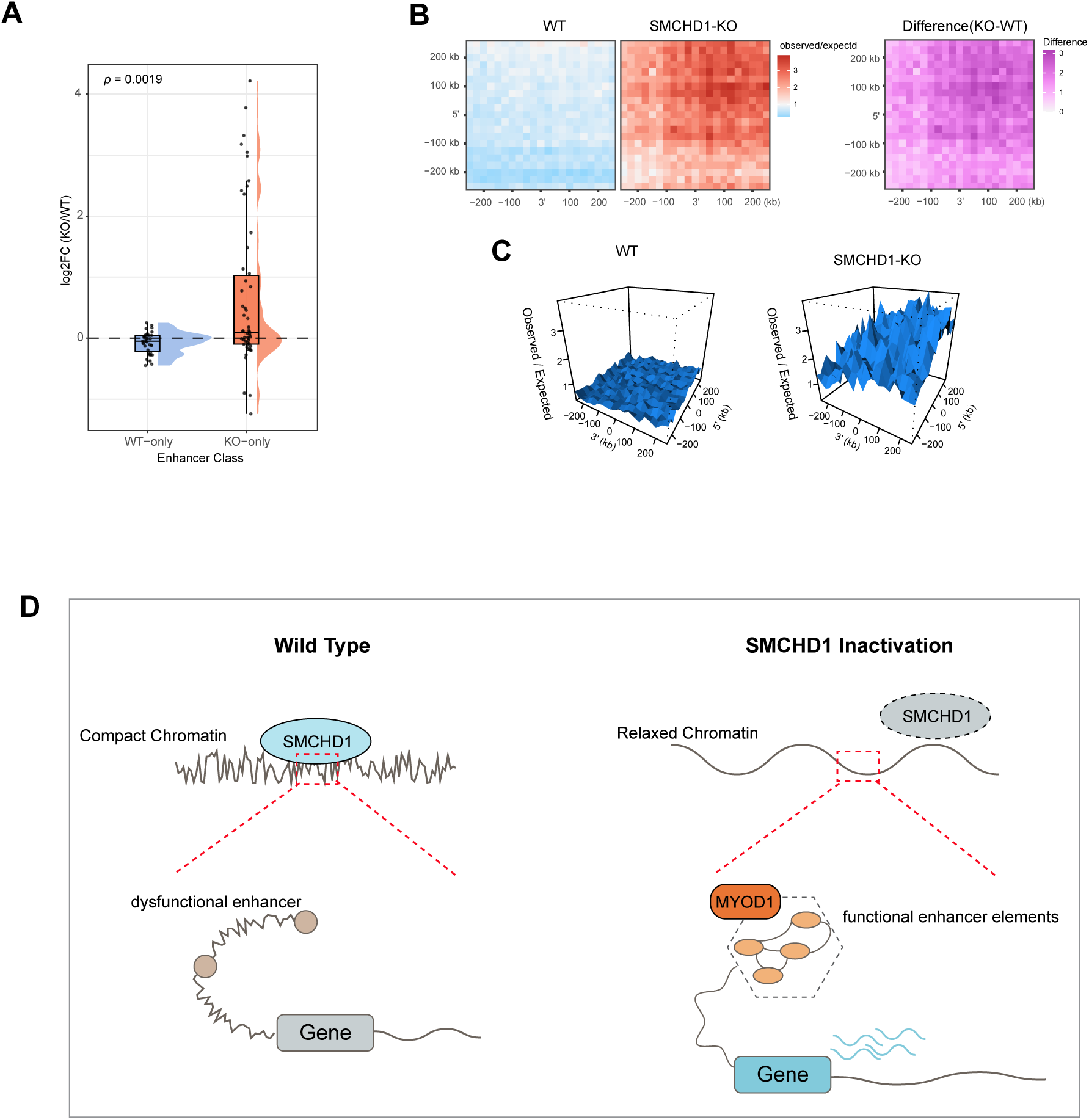
SMCHD1 inactivation increases local 3D chromatin contacts around MYOD1 enhancer nexus. **A.** Rain-cloud plot showing the log2 fold change of ATAC-seq signal in all enhancer elements within MYOD1 enhancer nexuses. The midline is the median of the data, with the upper and lower limits of the box being the third and first quartile (75th and 25th percentile), respectively. Whiskers extend to the most extreme data points within 1.5 × inter-quartile range (IQR) from the box bounds, and minima and maxima within the whiskers indicate the range of non-outlier data. Statistical significance was assessed using a two-sided Wilcoxon rank-sum test (*p* = 0.0019). **B.** 2D contacts heatmaps computed with GENOVA showing observed/expected (O/E) Hi-C interaction frequencies surrounding the centers of all enhancer elements. Left: WT cells show a relatively weak and diffuse contact pattern surrounding enhancer elements. Middle: SMCHD1-KO cells exhibit globally elevated O/E contacts, indicating increased local chromatin interaction frequency centered on enhancer elements. Right: The difference heatmap (KO − WT) highlights widespread gain of contact intensity within ±200 kb of enhancer elements in SMCHD1-KO cells, consistent with SE-proximal chromatin becoming more interactive after loss of SMCHD1. **C.** 3D surface plot showing the topology of contact enrichment around SE anchors. WT cells exhibit a relatively flat landscape, whereas SMCHD1-KO cells show a pronounced increase in local peaks, indicating a higher number of contacts over SEs. **D.** Model diagram for how SMCHD1 governs the MYOD1 enhancer nexus network.

## DISCUSSION

In this study, we present a previously unproven chromatin architectural role for SMCHD1 in restraining enhancer-driven transcriptional programs in human myoblasts. By integrating transcriptomic, epigenomic, and 3D genome contact analysis, we show that SMCHD1 inactivation leads to muscle cell-relevant transcriptional dysregulation that is independent of canonical DUX4-associated targets. Moreover, SMCHD1 loss results in global increases in chromatin accessibility, reorganization of higher-order chromatin architecture, and redistribution of the lineage-defining transcription factor MYOD1. Together, our results identify SMCHD1 as a key regulator of chromatin architecture that preserves transcriptional balance by limiting enhancer accessibility, enhancer clustering, and enhancer network– gene communication in the myogenic system.

Our data support a model in which SMCHD1 functions as a chromatin architectural regulator, rather than a locus-specific transcriptional regulator. In wild-type myoblasts, SMCHD1 maintains a relatively compact chromatin state at potential enhancers, restricting excessive chromatin opening and limiting local self-interaction. SMCHD1 loss relieves this compact chromatin state, leading to increased chromatin accessibility and increased enhancer activity. These results align with prior work showing that SMCHD1 contributes to chromatin compaction and structural regulation, such as inactive X chromosome compaction and the architecture at some autosomal loci ^8,61^. These effects are particularly pronounced at enhancer elements associated with MYOD1, indicating that SMCHD1 plays a critical role in modulating the responsiveness of lineage-specific regulatory programs to alterations in chromatin state.

A central finding of this work is the identification of MYOD1-related enhancer networks, which we termed MYOD1 enhancer nexuses. They are characterized by coordinated activation of multiple enhancer elements, including super-enhancers and stitched enhancers defined by ROSE (Figure 5), modulating target genes through reinforced chromatin looping, with locus-level examples (*CCL2* and *FAM155A*) (Figure 6). Unlike isolated enhancer–promoter interactions, a MYOD1 enhancer nexus represents higher-order regulatory units, including multiple enhancers, that are structurally and functionally integrated (Figure 6). This nexus-level architecture is consistent with the broader concept that enhancers can form cooperative multi-enhancer interaction networks, as demonstrated by multiple chromatin conformation approaches and high-resolution 3D mapping^62,63^. Our loop chaining and ABC modeling demonstrated that increased enhancer-gene contact activity within these nexuses is strongly associated with transcriptional activation, supporting a direct link between enhancer rewiring and gene activation in the absence of SMCHD1.

The formation of MYOD1 enhancer nexuses was characterized by increased chromatin accessibility and enhanced 3D chromatin interactions (Fig. 7B-C). Hi-C analysis revealed that SMCHD1 removal increases local contacts surrounding enhancer elements, resulting in more highly interactive enhancer elements (Fig. 7B-C). Three-dimensional representations of these contacts highlight the emergence of prominent interaction peaks in knockout cells, consistent with excessive enhancer self-connectivity (Fig. 7B-C). These findings suggest that SMCHD1 normally suppresses the formation of hyper-interactive enhancer networks, thereby preventing enhancer-gene communication. This observation is consistent with known principles that 3D genome structure both facilitates and constrains enhancer-promoter communication, and that enhancer–promoter proximity dynamics are functionally relevant for transcriptional expression^57,64^.

Importantly, an enhancer nexus arises preferentially from super- or stitched enhancers (Fig. 5, Fig. 6A), indicating that SMCHD1 loss promotes the activation of regulatory elements in the nexus. Furthermore, this finding is consistent with the concept that robust transcription can be driven by the coordinated activity of clustered enhancers, originally defined as super-enhancers, enriched for transcriptional machinery and master regulators, and linked to strong, identity-defining transcriptional programs^29,65^. SMCHD1 KO-specific enhancers within these nexuses underscore the role of SMCHD1 in safeguarding the enhancer landscape against inappropriate activation. By constraining the accessibility and interaction of chromatin, SMCHD1 limits MYOD1’s ability to engage in extensive enhancer clustering, thereby restricting transcriptional activation. Therefore, MYOD1 redistribution (Figure 4) suggests that SMCHD1 loss creates a more permissive chromatin state that expands or redirects MYOD1 occupancy, which in turn promotes enhancer activation and nexus formation (Figures 5–6). This is supported by prior work, which has demonstrated that MYOD1 binds widely across the genome in myogenic cells and is associated with broad changes in chromatin state, supporting its capacity to drive global regulatory remodeling when chromatin constraints are relaxed^48^. Taken together, our finding is also supported by the idea, which has illustrated that super-enhancers can concentrate coactivators and transcriptional apparatus via condensation-like behavior^66^, providing a plausible framework for hyper-connectivity of enhancer networks activating transcription once structural constraints are removed, such as those imposed by SMCHD1.

Our findings also provide a broader view for understanding transcriptional dysregulation mediated by SMCHD1 deficiency beyond DUX4-centered models (Fig. 1). While it is widely accepted that aberrant DUX4 expression remains an important pathogenic molecular mechanism in FSHD^32,34,67,68^, a number of recent studies have found that *DUX4* expression is hardly detectable in FSHD patient cells^35,39-41^, suggesting the existence of additional pathogenic mechanisms in FSHD. More importantly, our results demonstrate that SMCHD1 loss alone is sufficient to reprogram the epigenome landscape and chromatin architecture and may activate possible lineage-unrelated enhancer networks. This suggests that SMCHD1 deficiency creates a permissive chromatin environment that promotes transcriptional activation driven by transcription factors such as MYOD1, potentially contributing to disease heterogeneity and early pathogenic changes.

Our study extends the understanding of the function of SMCHD1 as a chromatin structural regulator. SMCHD1 not only shares functional similarities with other architectural regulators such as CTCF, cohesion, and the SMC protein family, but also serves a distinct role in the communication between the enhancer network and gene expression. Like CTCF, cohesin and SMC protein complexes primarily shape the 3D chromatin structure, including topologically associating domains, and loop boundaries^69-75^. SMCHD1 also selectively reshapes the 3D genome architecture of MYOD1-related enhancer-networks, making SMCHD1 a distinct modulator of lineage-specific enhancer network formation.

### Limitations of the study

Several limitations of this study should be acknowledged. First, our study was performed in a human immortalized primary myoblast model, and further validation in *in vivo* systems will be necessary to establish the generality of these mechanisms. Second, while ABC modeling, Hi-C and loop-chaining provide strong correlative evidence linking enhancer-network rewiring to transcriptional activation, direct functional perturbation of individual enhancer nexuses will be a future research direction. Third, while we obtained the high-resolution Hi-C contacts data for the whole genome, some loci were still not covered, resulting in missing connectivity of enhancers and gene expression, such as for the *FRG2B* gene, which is a potential FSHD2-associated gene found in our study (Figure S5B). Finally, our analysis captures a static snapshot following SMCHD1 loss, and the temporal dynamics of enhancer nexus formation and stabilization remain to be explored in the future.

Despite these limitations, our work establishes a conceptual framework in which SMCHD1 functions as a critical chromatin architectural safeguard that limits enhancer nexus formation and transcriptional activation. By restraining chromatin accessibility and local 3D chromatin interactions, SMCHD1 preserves regulatory stability in myoblasts and prevents excessive activation of MYOD1-involved enhancer network-gene programs. More broadly, these findings suggest that disruption of architectural repression can activate latent enhancer networks, providing a general mechanism by which chromatin regulators shape lineage-specific transcriptional outcomes in development and disease.

## RESOURCE AVAILABILITY

### Lead contact

Further information and requests for resources and reagents should be directed to and will be fulfilled by the lead contact, Gerd Pfeifer.

### Materials availability

All plasmids and cell lines generated in this study are available upon reasonable request from the lead contact.

### Data and code availability

The RNA-seq data are available in the GEO database (accession number GSE319423, reviewer token ufqhwqqqvvupxml). The ATAC-seq data can be found in the GEO database (accession number GSE319462, reviewer token qxinukuutdaddqv). The MYOD1 ChIP-seq data has been deposited into the GEO database (accession number GSE319422, reviewer token ihwbkmykhnkdvgv).

The loop chaining script is available here: (https://github.com/knight-hzj/published_script/tree/main/Pfeifer_lab/loop_chanining).

## ACKNOWLEDGEMENTS

We are indebted to the Shared Resource Facilities of the Van Andel Institute. This work was supported by grant R01AR079174 from the National Institute of Arthritis and Musculoskeletal and Skin Diseases (NIAMS) to G.P.P. The content is solely the responsibility of the authors and does not necessarily represent the official views of the National Institutes of Health.

## AUTHOR CONTRIBUTIONS

Conceptualization, Z.H. and G.P.P.; methodology, Z.H. and W.C.; investigation, Z.H., W.C., A.K. and G.P.P.; writing – original draft, Z.H.; writing – review & editing, Z.H., W.C., A.K. and G.P.P..; funding acquisition, G.P.P.; supervision, G.P.P.

## DECLARATION OF GENERATIVE AI AND AI-ASSISTED TECHNOLOGIES IN THE WRITING PROCESS

During the preparation of this manuscript, the authors used ChatGPT to improve readability in portions of this manuscript and Grammarly for spelling and grammar. After using this tool/service, the authors reviewed and edited the content as needed and take full responsibility for the content of the publication.

## METHODS

### Cell lines

LHCN-M2, a human male myoblast cell line immortalized with hTERT and CDK4, was obtained from Evercyte (Vienna, Austria). LHCN-M2 cells were cultured on 0.1% gelatin-coated tissue culture plates in DMEM / medium 199 (4:1) supplemented with 15% fetal bovine serum, 0.02 M HEPES, pH 7.1, 0.03 μg/mL zinc sulfate, 1.4 μg/mL vitamin B12, 0.055 μg/mL dexamethasone, 2.5 ng/mL hepatocyte growth factor, 10 ng/mL basic FGF, 60 units/mL penicillin and 60 μg/mL streptomycin.

### Generation of knockout LHCN-M2 myoblast lines using CRISPR-CasG

SMCHD1^-/-^ LHCN-M2 cell lines were generated by pLentiCRISPR-E with the blasticidin resistance gene generated from pLentiCRISPR-E-puromycin (a gift from Phillip Abbosh; Addgene plasmid # 78852), as in our previous study^10^. The lentiviral vector carrying SMCHD1 sgRNAs was co-transfected into HEK293FT cells with lentiviral packing plasmid psPAX2 and envelope plasmid pMD2.GVG to get viral particles. LHCN-M2 cells were transduced with plentiCRISPR-E-Blast-SMCHD1 virus, and single-cell clones were selected in 10 μg/mL blasticidin. Protein expression levels in the knockout cells were confirmed by Western blot. The genotype of all SMCHD1 knockout clones was further confirmed by the Sanger sequencing of the CRISPR-targeted region, which was PCR-amplified from genomic DNA and cloned into Topo-TA cloning vector (Thermo, 450030) as in our previous study^10^.

### RNA preparation and total RNA sequencing

We followed our standard procedures from a previous study^76^. The total RNA was purified from LHCN-M2 cells using the PureLink RNA Mini Kit (Ambion, 12183020) according to the manufacturer’s protocol. Total RNA QC was verified by Agilent 2100 Bioanalyzer (Agilent Technologies) and quantified with a NanoDrop 8000 instrument (Thermo Fisher). RNA-seq libraries were prepared from total RNA with the KAPA RNA HyperPrep Kit (KAPA Biosystems, KR1351) according to the manufacturer’s protocols. RNA-seq was performed with three biological replicates (for both wildtype and SMCHD1 knockout clones). The size distributions of the library were then validated on the Bioanalyzer (Agilent Technologies). Sequencing was performed with an Illumina NextSeq500 system. Library de-multiplexing was performed following Illumina standards.

### ATAC-seq

To map the profile of chromatin accessibility, we generated ATAC-seq libraries using the ATAC-seq Kit (Active Motif, 53150) according to the manufacturer’s protocol. Briefly, 100,000 fresh cells were centrifuged at 500xg for 5 minutes at 4°C, the supernatant was removed by pipetting, and 100 μl of ice-cold PBS was added, followed by centrifuging once more at 500xg for 5 minutes at 4°C. The supernatant was then removed, followed by resuspending the cell pellet in 100 μl ice-cold ATAC lysis buffer. Then, the resuspended cell pellet was centrifuged immediately at 500xg for 10 minutes at 4°C. Each lysed cell pellet was treated with 50 μl of Tagmentation Master Mix (2X tagmentation buffer, 10X PBS, 1.0% digitonin, 10 μl assembled transposomes) and suspended with gentle pipetting and then incubated at 37°C for 30 minutes in a thermomixer at 800 rpm. Following the tagmentation reaction, we transferred each sample to a new 1.5 ml microcentrifuge tube and added 250 μl DNA Purification Binding Buffer and 5 μl 3 M sodium acetate. We transferred the sample into the column and centrifuged at 17,000xg for one minute. We washed the column with 80% ethanol, then eluted the DNA from the column with 35 μl elution buffer. Library PCR was performed using the following program on a thermal cycler (72°C 5 minutes, 98°C for 30 seconds, 10 cycles of: 98°C for 10 seconds, 63°C for 30 seconds, 72°C for 1 minute, then hold at 10°C). We performed SPRI beads clean-up with 60 μl SPRI bead solution according to the manufacturer’s protocol, eluting the library DNA in 20 μl DNA Purification Elution Buffer. Sequencing was performed with an AVITI instrument with 150-bp paired-end read runs. All ATAC-Seq experiments were performed in three parallel wild-type and knockout samples.

### ChIP-seq

The binding profiles for MYOD1 protein were generated by ChIP-seq. We followed our standard procedures described in a previous study^10^. Briefly, cells were fixed with 1% formaldehyde for 10 min at room temperature and the crosslinking was quenched with 125 mM glycine. Fixed cells were then lysed with lysis buffer (50 mM HEPES-KOH, pH 7.9, 140 mM NaCl, 1 mM EDTA, 10% glycerol, 0.5% NP40, 0.25% Triton X-100), supplemented with cOmplete protease inhibitors (Roche)) and incubated on ice for 10 min. The lysed cells were then centrifuged at 3000xg for 5 min at 4°C in a benchtop centrifuge, followed by washing with wash buffer (10 mM Tris-Cl, pH 8.1, 200 mM NaCl, 1 mM EDTA, pH 8.0, 0.5 mM EGTA, pH 8.0) supplemented with cOmplete proteinase inhibitors and shearing buffer (0.1% SDS, 1 mM EDTA, 10 mM Tris-Cl, pH 8.1) supplemented with cOmplete proteinase inhibitors. Washed cell pellets were then resuspended in shearing buffer and sonicated with a Covaris E220 Evo sonicator to shear the DNA to 300 to 500 bp size fragments. The sheared lysate was supplied with NaCl and Triton X-100 to reach a final concentration of 150 mM NaCl and 1% Triton X-100, then cleared by centrifugation for 10 min at 20,000xg, and then incubated with washed Dynabeads Protein G (Invitrogen, 10004D) and MYDO1 antibody (Abcam, ab133627), overnight on a rotator at 4°C.

Beads were then collected and washed using low salt buffer (0.1% SDS, 1% Triton X-100, 2 mM EDTA, 20 mM HEPES-KOH, pH 7.9, 150 mM NaCl) supplemented with cOmplete proteinase inhibitors, high salt buffer (0.1% SDS, 1% Triton X-100, 2 mM EDTA, 20 mM HEPES-KOH, pH 7.9, 500 mM NaCl) supplemented with cOmplete proteinase inhibitors and LiCl buffer (100 mM Tris-Cl, pH 7.5, 500 mM LiCl, 1% NP40, 1% sodium deoxycholate) twice and with TE buffer (10 M Tris-Cl, pH 8.1, 1 mM EDTA) once. ChIP DNA samples were eluted with Proteinase K digestion buffer (20 mM HEPES, pH 7.9, 1 mM EDTA, 0.5% SDS) and 1 µl of Proteinase K (20 mg/ml). Purified DNA was quantified for library preparation with Qubit sensitivity dsDNA HS Kit. Libraries were then prepared using the TruSeq ChIP Sample Preparation Kit (Illumina, IP-202–1012, IP-202–1024) according to the manufacturer’s instruction. Briefly, 5 ng of ChIP-DNA was used for input and IP samples. Libraries were amplified using 14 cycles on a thermocycler. Libraries were then quantified and validated using the Agilent High Sensitivity DNA Kit and bioanalyzer. Sequencing was performed with an Illumina HiSeq 2500 instrument with 150-bp paired-end read runs. The ChIP-Seq experiments were processed in parallel with whole cell extract input controls.

### Quantification and Statistical Analysis

#### RNA-seq data analysis

For RNA-seq analysis, we followed our standard procedures^76^. Seventy-five base pair reads were trimmed with Trim Galore (v0.4.0); trimmed reads were then aligned to the human genome hg19 with STAR (version 2.5.1), and gene count was calculated with STAR^77^. Differential gene expression was called by the Limma (v3.38.2) statistical package^78^. P-values for differential expression were adjusted for multiple testing correction using the Benjamini-Hochberg method in the stats package. Statistical significance for differentially expressed genes was fold change >2 and q <0.05. Differential gene expression was plotted with the R package ggplot2 (v3.5.2) and pheatmap (v1.0.12). The GO function annotation was performed with DAVID^36-38^.

#### ChIPseq data analysis

The adapter and low-quality sequences were trimmed from 3′ and 5′ ends by Trimgalore (v0.5.0). Subsequently, the trimmed reads were aligned to hg19 with Bowtie2 (versionv2.5.1). The aligned reads were then deduplicated using PicardTools (http://broadinstitute.github.io/picard). ChIP-peaks were then analyzed with MACS2^79^. All ChIP data was normalized against the corresponding input controls using the ‘-c’ option of MACS2. For MYOD1, ChIP-Seq peaks were called using the ‘callpeak’ function of MACS2 with --nomodel -f BAMPE -g hs -q 0.05 -B. For H3K27ac, H3K4me1, we used the peak information from our previous study^10^. Bigwig tracks of ChIP-seq were generated using bamCoverage of deeptools^80^ (v3.5.5) to compute coverage across the entire genome. Heatmaps and enrichment profiles of average Counts Per Million mapped reads (CPM) were performed using deeptools. Overlap of genomic regions and peaks contained within them was determined with bedtools (v2.30.0). The enrichment profile plot of MYOD1 was generated with deeptools. Differential MYOD1 peak binding was calculated with R package csaw^81^ (v1.28.0) and edgeR^82^ (v3.36.0).

#### Hi-C data analysis

For Hi-C data analysis, we followed our standard procedures^10^. The in situ Hi-C data were processed with a standard pipeline of Juicer^83^, using a pipeline code available at (https://github.com/theaidenlab). Within sample normalization was performed using the Knight-Ruiz (KR) method^84^. Hi-C data used in this study was comprised of two sets of biological replicates for each WT and SMCHD1 knockouts, the two biological replicates for each genotype were merged to get higher resolution (average of 4.8 billion contacts) for further analysis^10^. The compartment, TAD and loop information was obtained from our previous study, with more detail^10^. The 2D contact map along with the chromatin accessibility and A/B compartments scores^10^ were visualized by Juicer^83^. To show the chromatin contacts along with chromatin accessibility in the TADs between WT and SMCHD1 knockouts, FAN-C^85^ (v 0.9.1) was used to generate the 2D contact heatmaps, insulation scores of TADs and ATAC-signals. Differential TAD boundaries identified by the R package TADcompare (v1.18.0)^86^ from our previous study^10^ in the B-to-A region were used for calculating the chromatin accessibility. To show the average contact count distribution of loops and their surroundings in compartment B-to-A switched regions between WT and SMCHD1 knockouts, we performed an aggregate peak analysis (APA) of loop calls using the R package GENOVA ^60^ (v1.0.0.9.98) at 10 kb resolution.

#### ATAC-seq data analysis

Adapters and low-quality bases were trimmed from paired-end reads by Trimgalore (v0.5.0). Subsequently, the trimmed reads were aligned to hg19 with Bowtie2 (v2.5.1). The aligned reads were then filtered to remove mitochondrial DNA reads with samtools, followed by deduplication using PicardTools (http://broadinstitute.github.io/picard). Then, the deduplicated reads with low quality were removed with deeptools, and overlapping genomic blacklist regions were filtered with deeptools. Filtered BAM files were used to generate normalized genome browser tracks (BigWig) for visualization in the IGV browser. Chromatin accessibility peaks were identified with MACS2^79^. ATAC-Seq peaks were called using the ‘callpeak’ function of MACS2 with -f BEDPE --nomodel --shift -37 --extsize 73 -g hs -B –broad --keep-dup all --cutoff-analysis. Differential accessibility analysis was performed with R package csaw^81^ (v1.28.0). A consensus peak set was generated by merging high-confidence peaks across all samples. The accessibility read counts within these peaks were then quantified. The normalized count matrices were then analyzed for differential accessibility between experimental samples using statistical R package edgeR^82^ (v3.36.0). Peaks with an adjusted *p*-value < 0.05 and a fold change > 2 were considered significantly differentially accessible. Significantly differentially accessible regions were annotated with nearby genes and genomic features using annotatePeaks.pl of HOMER^87^ (v44.11.1). Motif enrichment analysis was performed on differentially accessible regions to identify overrepresented transcription factor binding motifs using JASPER (v2026), providing insight into regulatory factors potentially driving differential chromatin accessibility. The differential chromatin accessibly quantification at compartment switching regions and newly formed TAD boundaries were calculated with ggplot2(v3.5.2) and ggpubr (v0.5.6).

#### Identification of super-enhancers and stitched enhancers

Super-enhancers and stitched enhancers were defined using the ROSE^29^ program (http://younglab.wi.mit.edu/super_enhancer_code.html). We first collected the peak sites of H3K27ac and H3K4me1 using MACS2^79^ peak caller from our previous study^10^, and then the defined peaks were transformed into GFF files to meet the criteria of input files of the ROSE program. We ran the ROSE with stitching distance option of 12.5 Kb and promoter-proximal regions were excluded by removing peaks within ±2.5 kb of annotated transcription start sites (TSSs) to focus analysis on distal regulatory elements for WT and SMCHD1KO samples, respectively. Super-enhancers and stitched enhancers identified by both H3K27ac and H3K4me1 ranking were used for further analysis.

#### Definition of MYOD1 enhancer nexuses

We defined MYOD1 enhancer nexuses computationally by integrating enhancer activity, MYOD1 binding, chromatin looping, and Activity-by-Contact (ABC) style enhancer–gene scores in WT and SMCHD1-KO myoblasts. First, MYOD1 enhancer nexuses were constructed from HiC-derived loops and enhancer annotations. Hi-C loops were identified using HiCCUPS (https://github.com/theaidenlab) and exported as merged BEDPE files for WT and SMCHD1-KO myoblasts from our previous study^10^. Loop anchors were expanded by ±5 kb prior to overlap analyses. TAD domains (at 10 kb resolution) were identified using Juicer (https://github.com/theaidenlab) as in our previous study^10^. Enhancer elements including stitched enhancers and super-enhancers called by ROSE^29^ were defined from H3K27ac/H3K4me1 peaks and assigned unique IDs (*se_id*). A loop chaining module was then built around each MYOD1-bound enhancer element, for every element that overlapped a MYOD1 ChIP–seq peak defined as direct enhancers. All enhancers connected by at least one HiCCUPS loop within the same TAD were iteratively added, yielding a set of direct and related enhancers sharing a common loop-connected neighborhood. Each neighborhood was assigned a unique *group_id,* defined as the genomic coordinate of the MYOD1-bound index enhancer. These loop-connected enhancer element sets constituted the structural candidates for MYOD1-loop nexuses in each condition. Secondly, we performed modified ABC-style enhancer-gene scoring in WT and SMCHD1-KO conditions. Promoters were then defined as ±2 kb around transcription start sites from TxDb.Hsapiens.UCSC.hg19.knownGene(https://bioconductor.org/packages/TxDb.Hsapien <u>s.UCSC.hg19.knownGene/</u>). Promoter pairs were retained only if they have the same loop connected (promoter in anchor A with enhancer element in anchor B, or vice versa). For each enhancer element–gene pair connected by at least one HiCCUPS loop, loop contact strength was quantified as observed/expectedBL from HiCCUPS output. A distance-weighted contact was computed using exponent α = 0.7, with a minimum distance of 5 kb. Enhancer activity (mean of H3K27ac and H3K4me1) was multiplied by distance-weighted contact to get *ABC_raw*. For each gene, ABC scores were normalized across enhancers within ±2 Mb of the TSS to get *ABC_norm*. Differential activity was computed as ΔABC = ABC_norm(KO) − ABC_norm(WT). ΔABC values were integrated with log₂fold changes and false discovery rates (FDRs) from RNAseq data. Thirdly, we quantified loop topology within MYOD1 enhancer nexuses using the enhancer–enhancer edge result derived from Hi-C loop chaining. For every nexus, we computed the number of super-enhancers and stitched enhancers, the maximum hop distance along the loop chain, and the fraction of adjacent pairs that were directly looped. Module closure was assessed as *first_last_direct,* indicating whether an enhancer element at the minimum hop layer and an enhancer element at the maximum hop layer were directly connected by an EE edge. The loop chaining script is available here: (https://github.com/knight-hzj/published_script/tree/main/Pfeifer_lab/loop_chanining).

---

**Supplementary Fig. 1:**
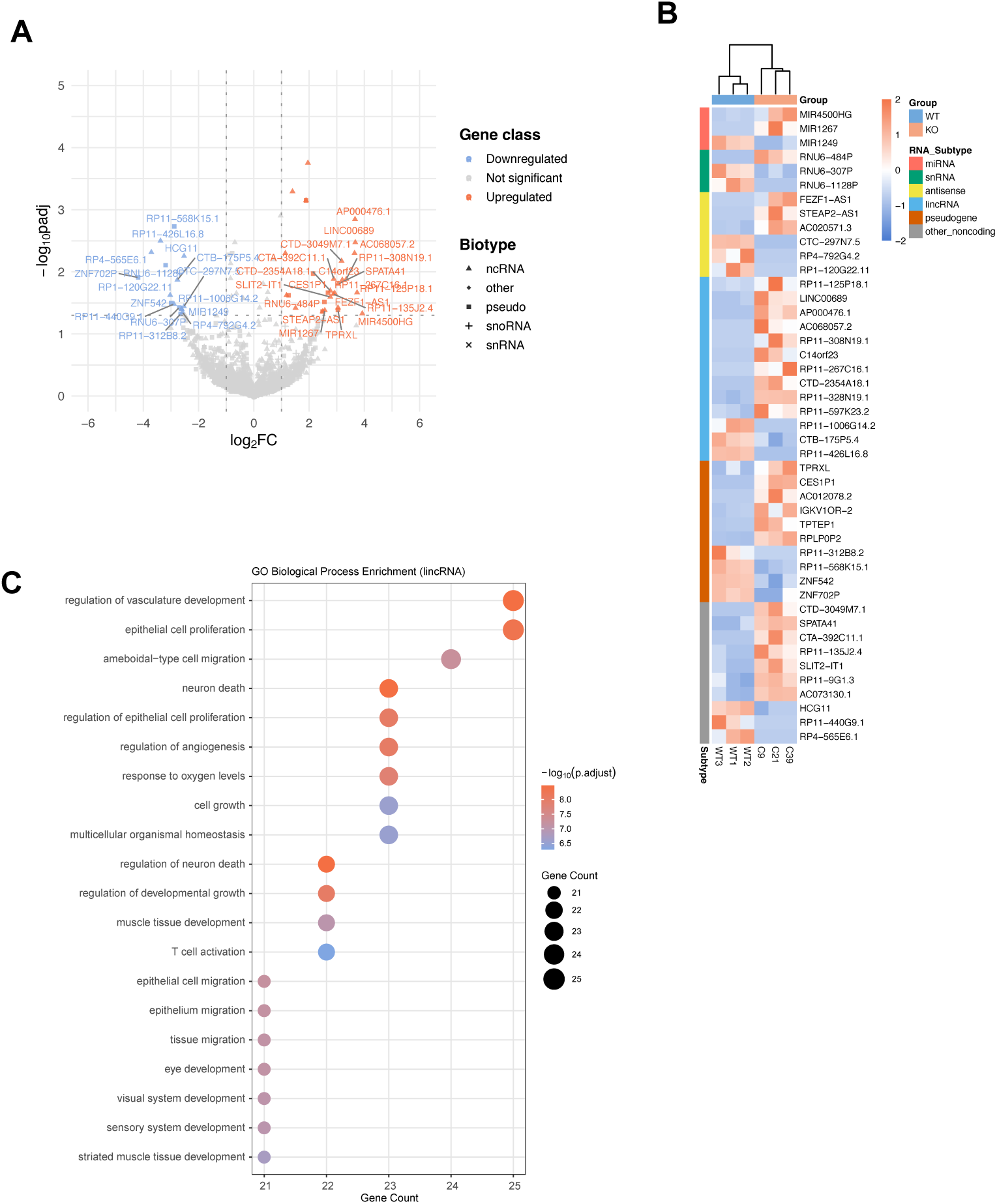
SMCHD1 loss leads to aberrant expression of non-coding RNAs. **A.** Volcano plot showing the expression of non-coding RNAs after SMCHD1 knockout. All differentially expressed non-coding RNAs with FDR <= 0.05 and log2 fold change >2 were labeled in light blue and red. Different symbols indicate the different non-coding RNA types. **B.** Heatmap showing the expression of all differentially expressed non-coding RNAs (from panel A) in three wildtype (WT) and three knockout myoblast clones. **C.** Scatter plot showing the gene functional analysis of differentially expressed lincRNAs.

**Supplementary Fig. 2:**
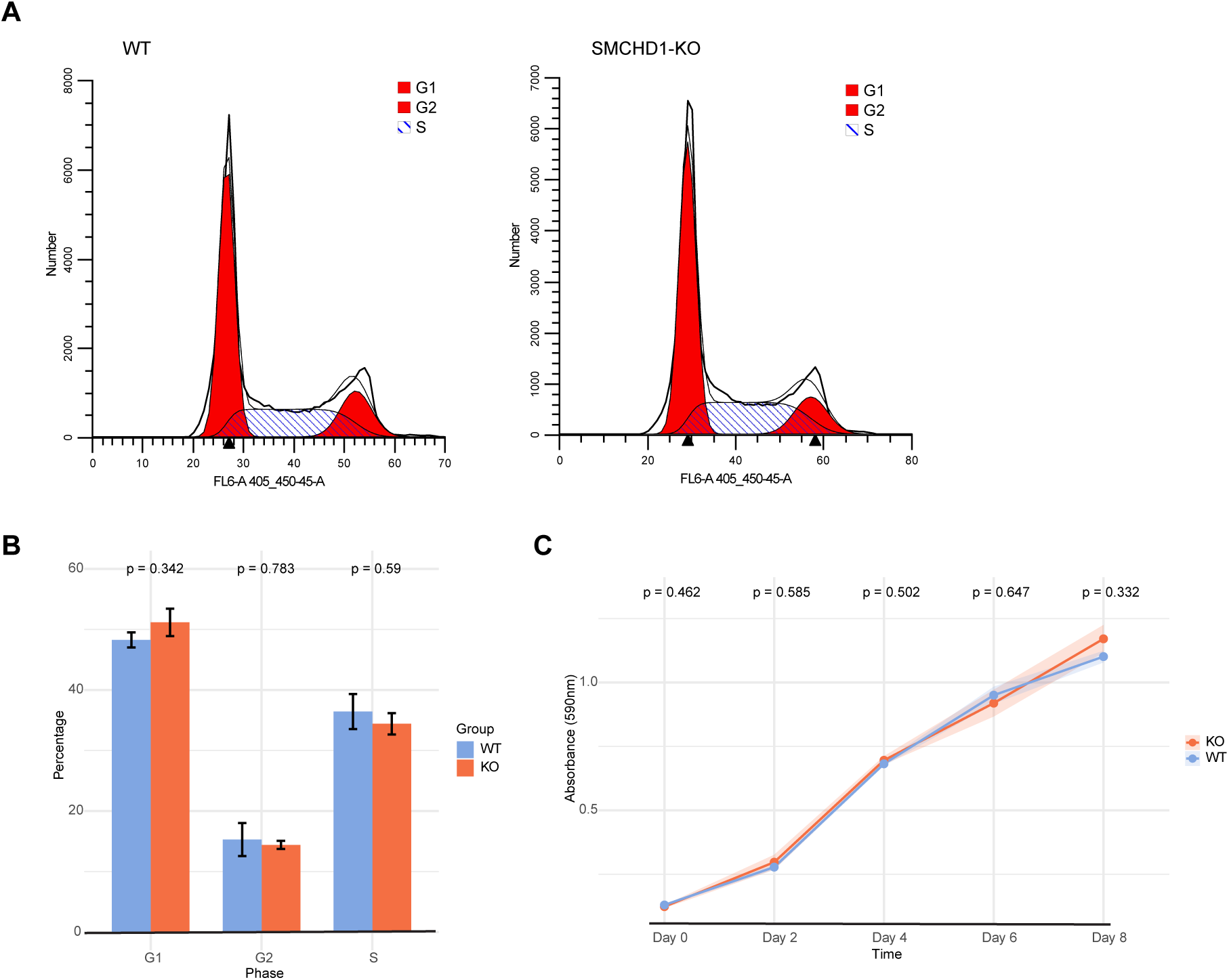
Cell cycle distribution and proliferation rate of LHCN-M2 wildtype and SMCHD1 knockout cells. **A.** Cell cycle profiles measured by FACS. **B.** Cell cycle phase distribution measured by FACS. Statistical significance was assessed using T-test. **C.** Cell growth rates were measured by MTT assay. Statistical significance was assessed using T-test.

**Supplementary Fig. 3:**
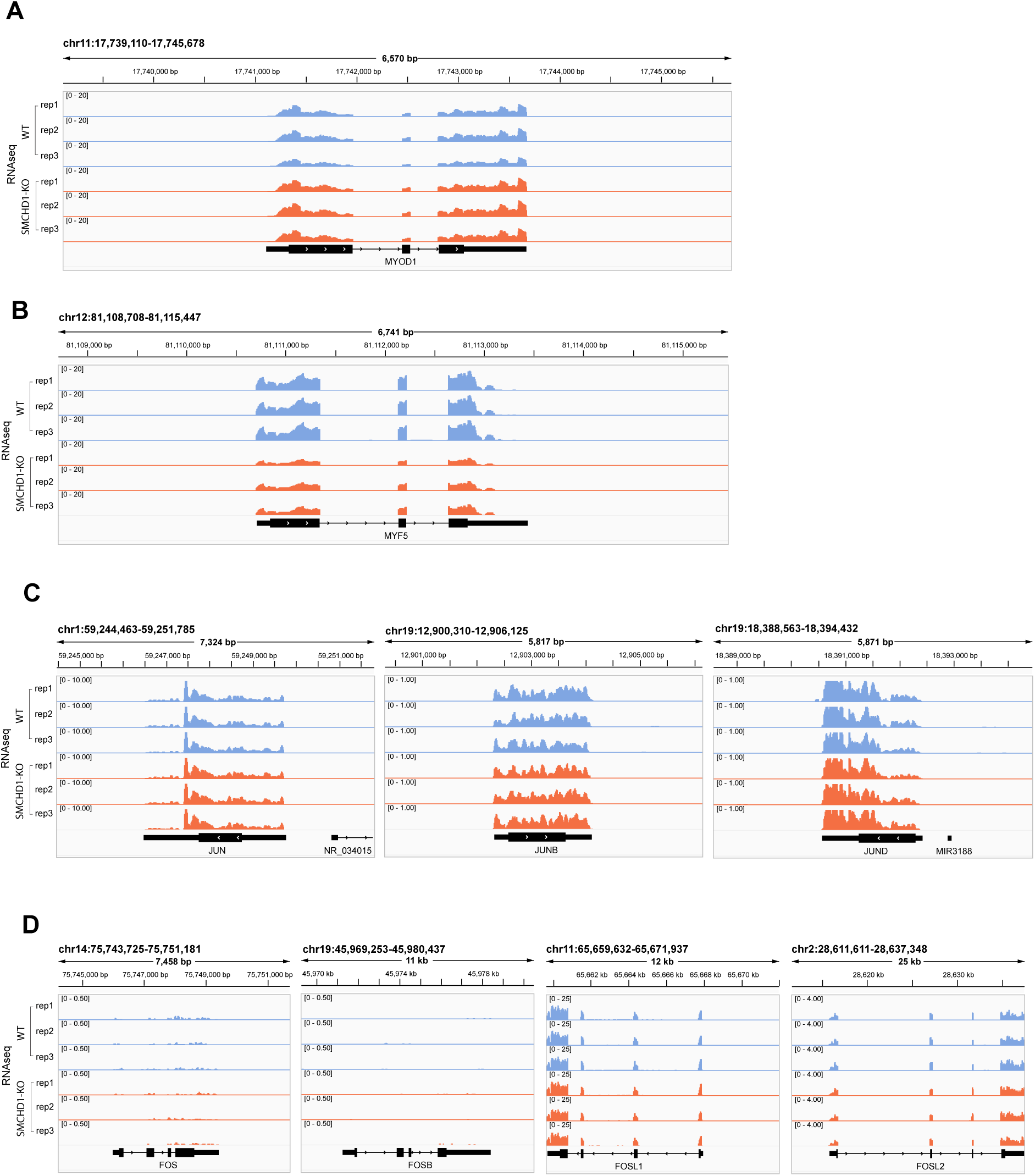
RNA expression level for *MYOD1*, *MYF5* and *JUN* family genes. **A.** Illustration of RNA-seq peaks at *MYOD1* gene locus. **B.** Illustration of RNA-seq peaks at *MYF5* gene locus. **C.** Illustration of RNA-seq peaks at *JUN* family genes loci. **D.** Illustration of RNA-seq peaks at *FOS* family genes loci.

**Supplementary Fig. 4:**
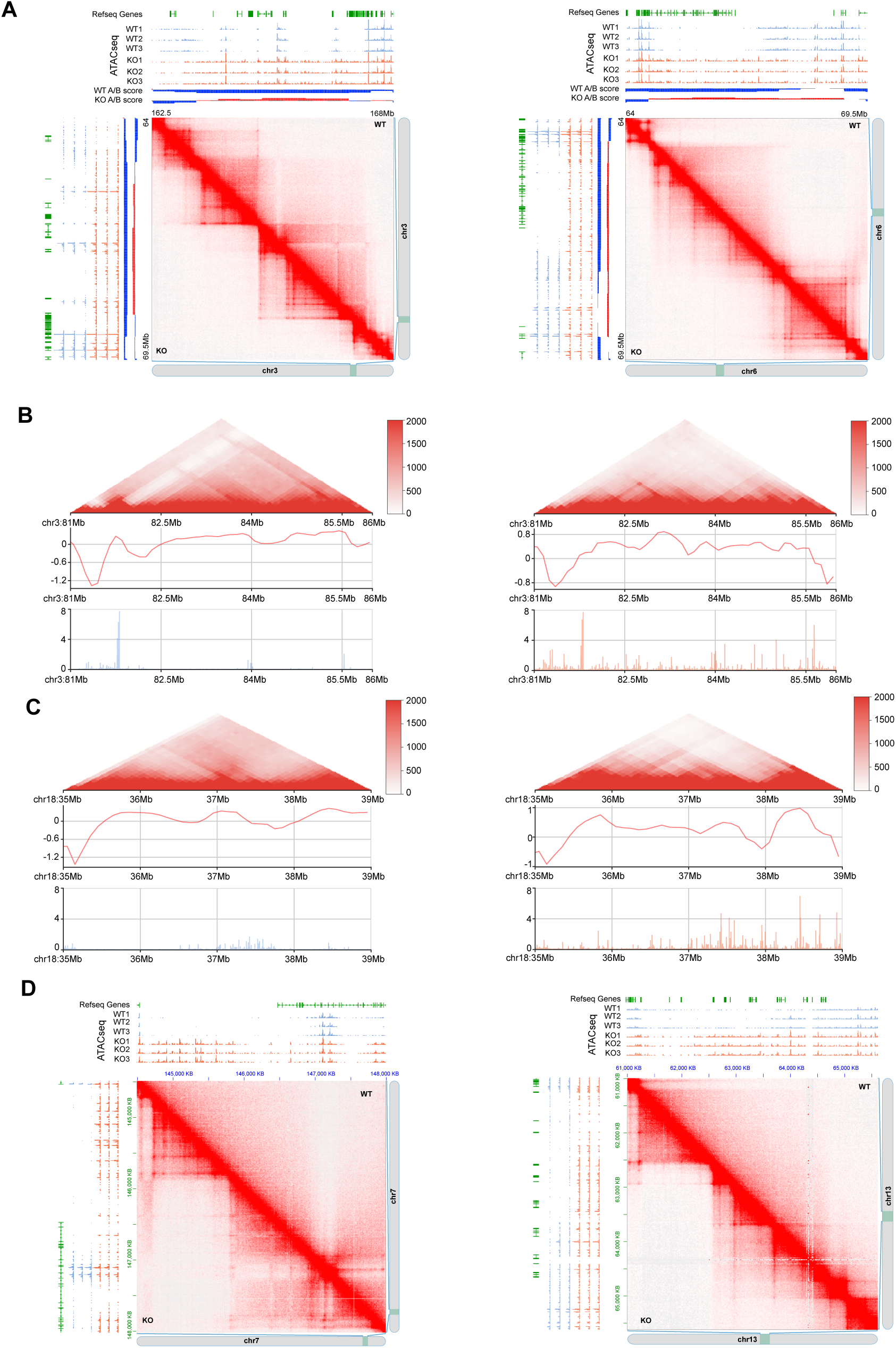
3D chromatin structure and accessibility in compartment switching regions after loss of SMCHD1. **A.** HiC contacts and Hi-C eigenvector (PC1) analysis in WT and SMCHD knockout cells across representative regions of chromosome 3 and 6. The ATAC-seq signals are shown along with B compartments (blue) and A compartments (red). B-to-A transitions can be seen in the knockout cells compared to the wildtype. **B-C.** Hi-C contact map, insulation profiles and ATAC-seq signal at representative regions on chromosome 3 and 18 in wildtype cells (left panel) and SMCHD1 knockout cells (right panel). **D.** Gains of loops at representative regions on chromosomes 7 (left panel) and 13 (right panel), linked to enhanced chromatin accessibility in SMCHD1 KO cells, as shown in the ATAC-seq tracks.

**Supplementary Fig. 5:**
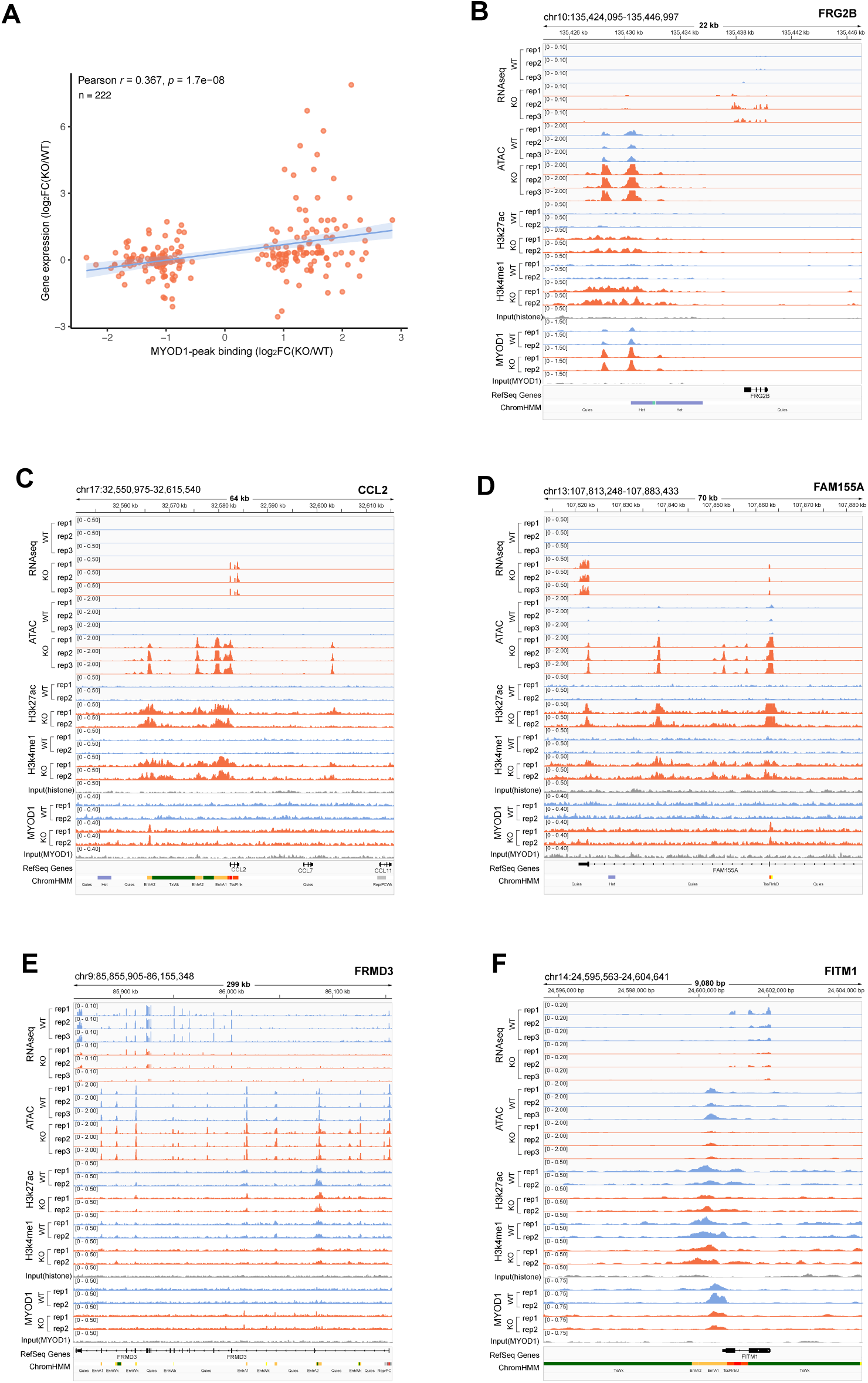
The relationship between MYOD1 binding and gene expression. **A.** Correlation between MYOD1 binding and altered gene expression. Pearson’s correlation coefficient was calculated with R to determine the strength and direction of the relationship between altered MYOD1 binding and altered gene expression. **B-D.** Illustration of RNA-seq peaks, ATAC-seq, H3K4me1, H3K27Ac, and MYOD1 ChIP seq at representative upregulated gene loci. **E-F.** Illustration of RNA-seq peaks, ATAC-seq, H3K4me1, H3K27Ac, and MYOD1 ChIP seq at representative downregulated gene loci.

**Supplementary Fig. 6:**
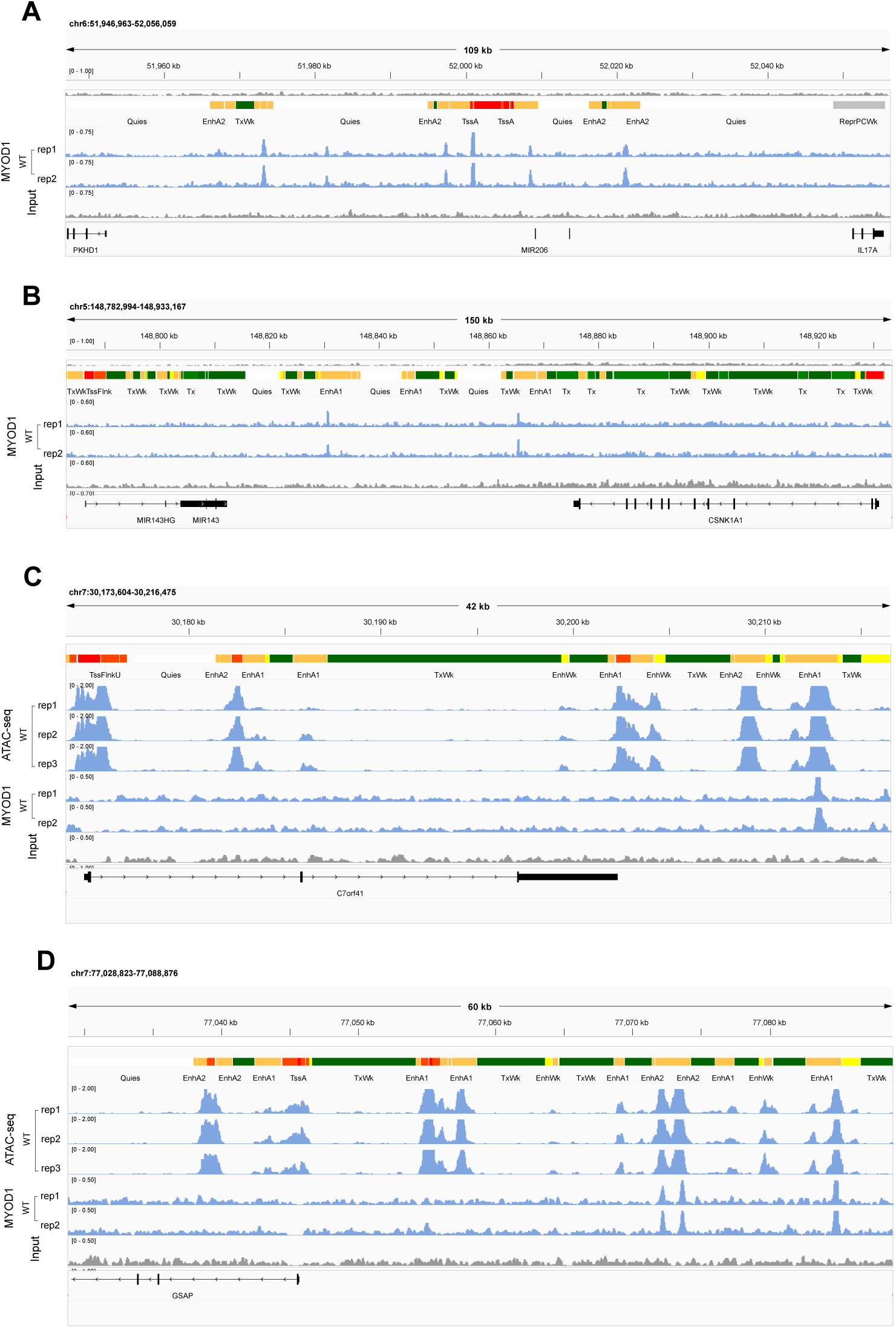
MYOD1 bound to the accessible enhancer regions. **A-B.** Illustration of IGV browser views showing MYOD1 localized to enhancer regions. **C-D.** Illustration of the IGV browser views showing MYOD1 preferentially localized to accessible enhancer regions.

**Supplementary Fig. 7:**
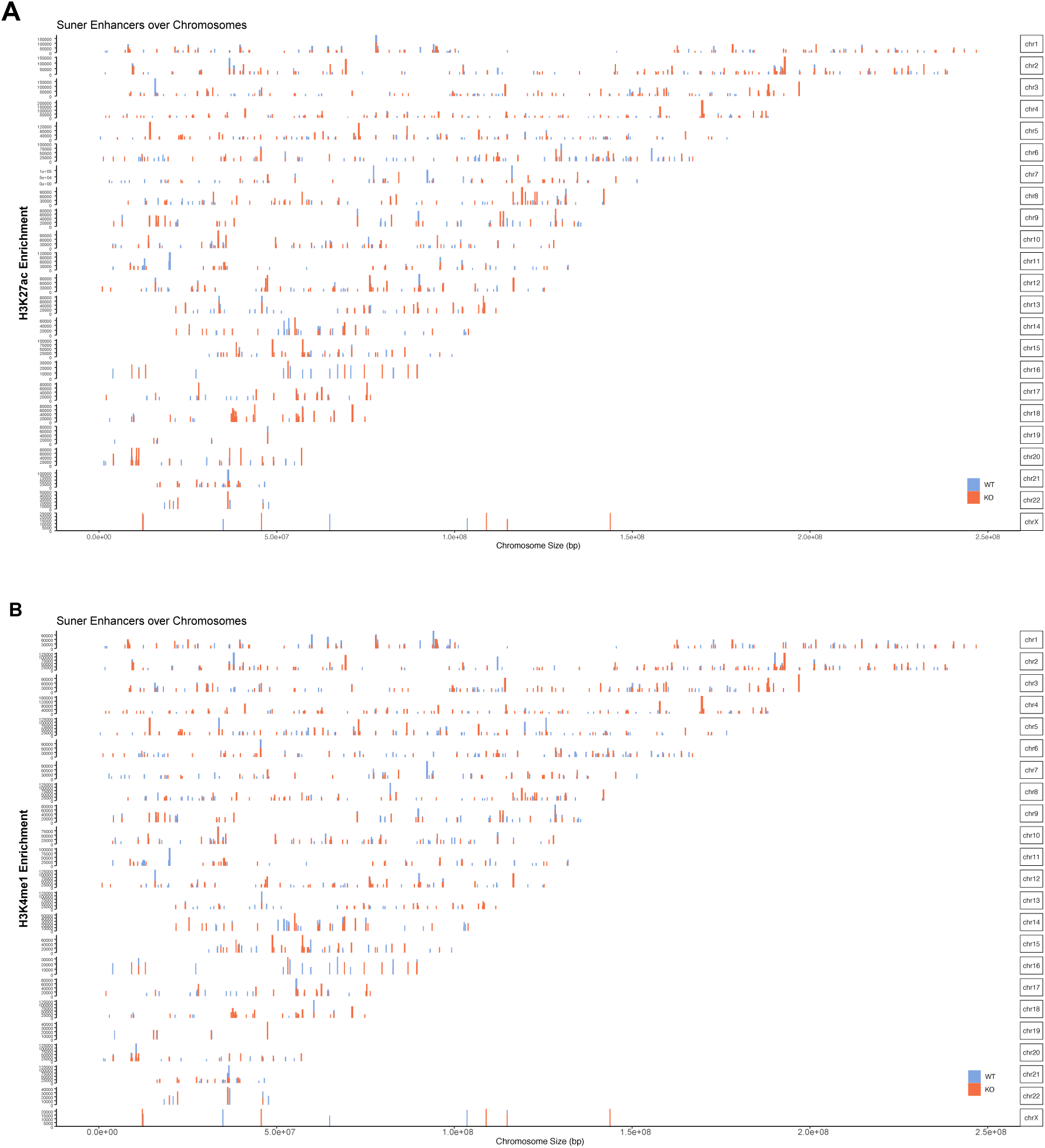
Distribution of super enhancers in the WT and SMCHD1 knockouts. **A.** H3K27ac enrichment pattern at super enhancers of WT and SMCHD1 knockouts. **B.** H3K4me1 enrichment pattern at super enhancers of WT and SMCHD1 knockouts.

**Supplementary Fig. 8:**
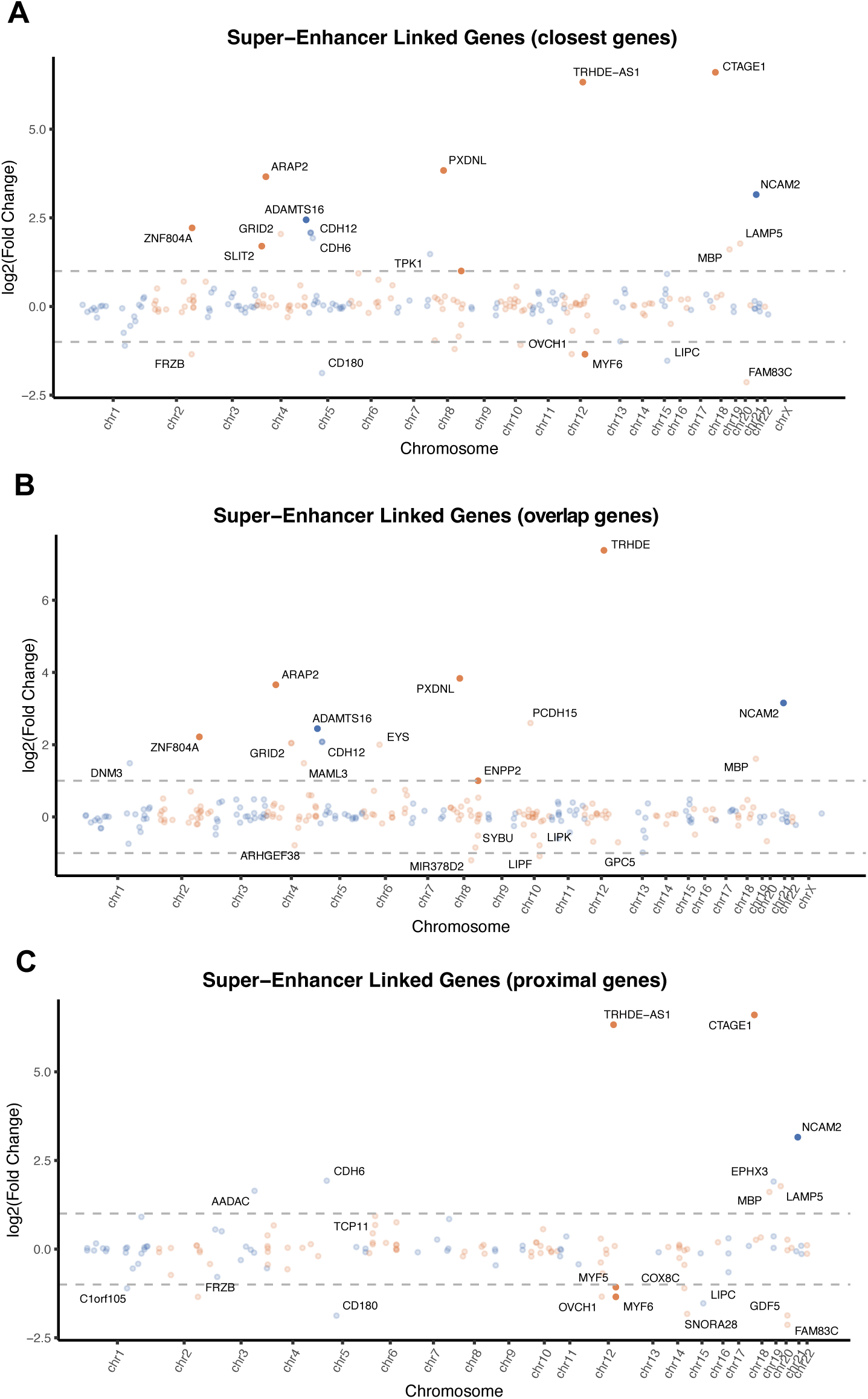
Super enhancer-linked genes that change expression in the absence of SMCHD1. **A.** Expression of super enhancer-closest genes (the single gene with the nearest TSS to the super enhancers). **B.** Expression of super enhancer-overlapped genes. **C.** Expression of super enhancer-proximal genes (genes within a fixed window of ±100kb from the SE coordinates).

**Supplementary Fig. 9:**
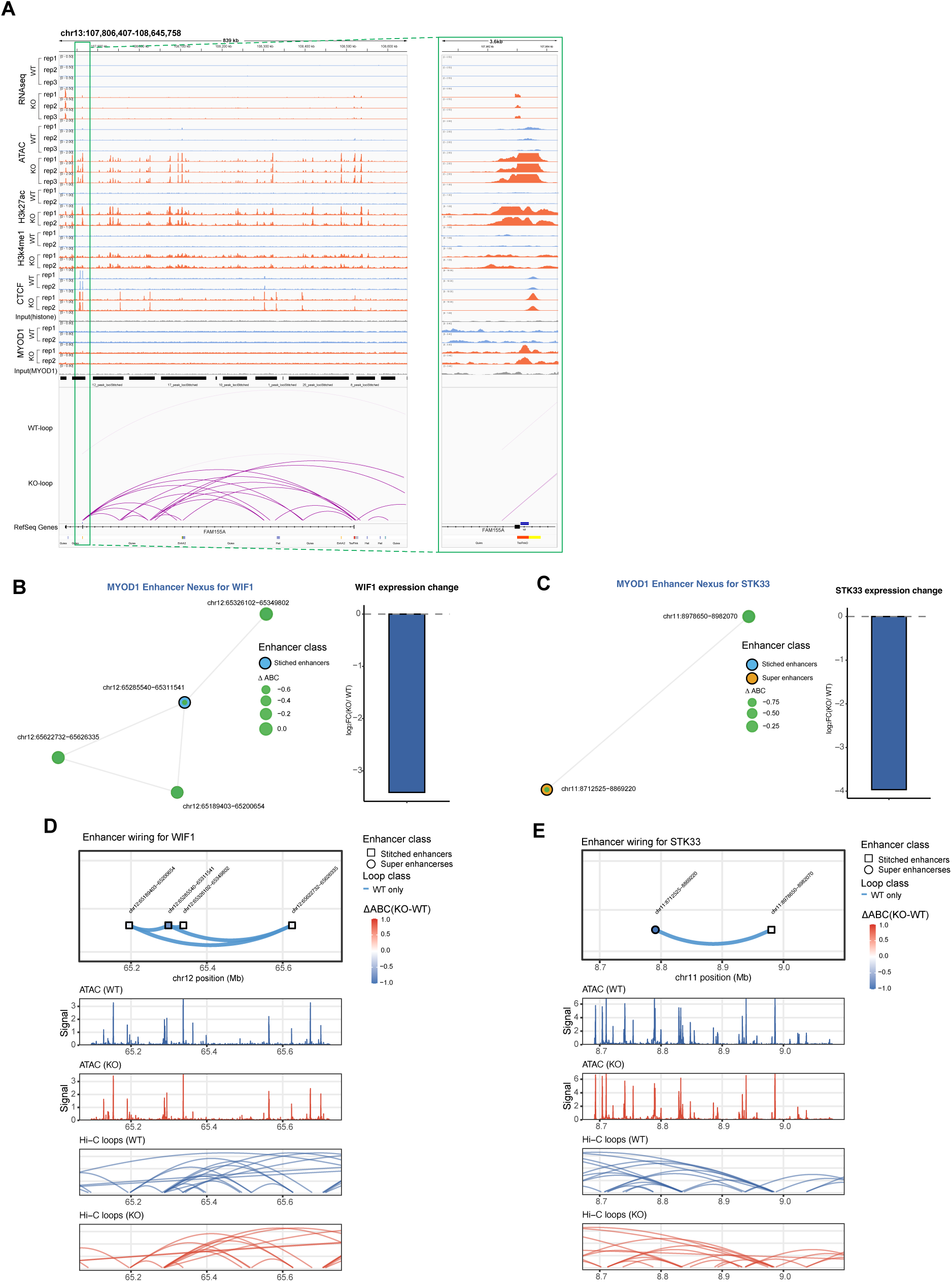
Transcriptional activation is mainly associated with the establishment of a MYOD1 enhancer nexus upon loss of SMCHD1. **A.** Illustration of the IGV browser showing *FAM155A* expression related to the more open chromatin accessibility, H3K27ac and H3K4me1 binding, and more loop contacts. The stitched enhancer called by ROSE was characterized by newly formed MYOD1 binding upon SMCHD1 loss. Enlargement of an area at the gene locus is shown in the green rectangle at the right side. **B.** MYOD1 enhancer nexus deactivated in SMCHD1 knockout for *WIF1*. Four WT-only enhancer clusters form a MYOD1enhancer nexus regulating *WIF1*, each showing enhancer activity and looping. Node size reflects enhancer ABC activity. *WIF1* expression exhibits a strong KO-induced decrease, as shown in the bar chart at the right. **C.** MYOD1 enhancer nexus deactivated in SMCHD1 knockout for *STK33*. Two WT-only enhancer clusters form a MYOD1 enhancer nexus regulating *STK33*, each showing enhancer activity and looping. Node size reflects enhancer ABC activity. *STK33* expression exhibits a strong KO-induced decrease, as shown in the bar chart at right. **D-E.** Enhancer wiring, peak tracks, and gene models for the representative MYOD1 enhancer nexuses for *WIF1* and *STK33*. Top: genomic enhancer wiring tracks show enhancer coordinates (black squares), KO-only enhancer arcs (orange), and ΔABC activity is defined by arc color. Middle: ATAC-seq signal confirms that WT and SMCHD1 KO have similar chromatin accessibility at enhancers. Bottom: Hi-C loop contact tracks illustrate WT and SMCHD1 KO have a similar chromatin contact pattern in the nexus.

